# Mystery of fatal ‘Staggering disease’ unravelled: Novel rustrela virus causes severe encephalomyelitis in domestic cats

**DOI:** 10.1101/2022.06.01.494454

**Authors:** Kaspar Matiasek, Florian Pfaff, Herbert Weissenböck, Claudia Wylezich, Jolanta Kolodziejek, Sofia Tengstrand, Frauke Ecke, Sina Nippert, Philip Starcky, Benedikt Litz, Jasmin Nessler, Peter Wohlsein, Christina Baumbach, Lars Mundhenk, Andrea Aebischer, Sven Reiche, Pia Weidinger, Karin M. Olofsson, Cecilia Rohdin, Christiane Weissenbacher-Lang, Julia Matt, Marco Rosati, Thomas Flegel, Birger Hörnfeldt, Dirk Höper, Rainer G. Ulrich, Norbert Nowotny, Martin Beer, Cecilia Ley, Dennis Rubbenstroth

**Author notes:** <u>Corresponding author:</u> Dennis Rubbenstroth.

## Abstract

‘Staggering disease’ is a neurological disorder considered a threat to European domestic cats (*Felis catus*) for almost five decades. However, its aetiology has remained obscure. Rustrela virus (RusV), a relative of rubella virus, has recently been shown to be associated with encephalitis in a broad range of mammalian hosts. Here, we report the detection of RusV RNA and antigen by metagenomic sequencing, RT-qPCR, *in-situ* hybridization and immunohistochemistry in brain tissues of 28 out of 29 cats with non-suppurative meningoencephalomyelitis and ‘staggering disease’-like neurological disorder from Sweden, Austria, and Germany, but not in non-affected control cats. Screening of possible reservoir hosts in Sweden revealed RusV infection in wood mice (*Apodemus sylvaticus*). Our work strongly indicates RusV as the long-sought cause of feline ‘staggering disease’. Given its broad host spectrum and considerable geographic range, RusV may be the aetiological agent of neuropathologies in further mammals, possibly even including humans.

## INTRODUCTION

Throughout mammalian species, inflammatory disorders of the central nervous system (CNS) are associated with substantial suffering, mortality and long-term neurological deficits. Aetiopathogenically, they can be broadly categorised into infectious and immune-mediated disorders^1^. All too often, however, the cause of an encephalitis remains unknown and leaves clinicians, patients and owners of affected pets with considerable uncertainty about its origin, treatment options and, hence, prognosis. The latter holds true particularly for the large histopathologically convergent panel of non-suppurative, lymphohistiocytic encephalitides. A considerable proportion of these cases remains unsolved using conventional diagnostic methods, such as immunohistochemistry (IHC), *in-situ* hybridization (ISH), and polymerase chain reaction (PCR) techniques for regional neurotropic pathogens^2, 3, 4, 5, 6^.

One of those controversial encephalitides of possibly infectious origin is the so-called ‘staggering disease’ of domestic cats (*Felis catus*). It has been described first in the 1970s in the Swedish Lake Mälaren region between Stockholm and Uppsala^7^, which remains a hotspot of ‘staggering disease’ to the present. Neurologic disorders possessing striking similarity with this disease entity have been described also in domestic cats in other European countries, particularly in Austria^6, 8, 9, 10, 11^, and even in other felids^12, 13^.

The most prototypic clinical sign is hind leg ataxia with a generally increased muscle tone resulting in a staggering gait. In addition, a broad range of other neurologic signs may occur, including the inability to retract the claws, hyperaesthesia and occasionally tremors and seizures. Behavioural alterations may range from enhanced vocalization, depression and becoming more affectionate to rarely aggression^7, 8, 14, 15^. The disease progression usually lasts a few days to a few weeks, but may also continue for more than a year, and it generally results in euthanasia. for animal welfare reasons. The histopathology of ‘staggering disease’ is characterized by a non-suppurative, predominantly lymphohistiocytic meningoencephalomyelitis with angiocentric immune cell invasion and perivascular cuffing predominantly in the grey matter of the CNS^7, 8, 14, 15, 16^.

While the microscopic pattern has suggested a viral origin of ‘staggering disease’, its aetiological agent has remained undisclosed for almost five decades. Borna disease virus 1 (BoDV-1; species *Orthobornavirus bornaense*; family *Bornaviridae*), which causes neurologic disorders in various mammals including humans^17^, has for a long time spearheaded the panel of aetiological candidates^16, 18, 19, 20, 21, 22, 23^. However, results suggesting natural BoDV-1 infections in cats with ‘staggering disease’ in Sweden remained inconclusive and were later refuted on the grounds of new standards^17, 24, 25^.

Fortunately, advances in clinical metagenomics over the last years have provided us with promising tools for the detection of new or unexpected pathogens involved in hitherto unexplained encephalitides^26, 27, 28, 29, 30, 31^. By application of an established metagenomic workflow^32^, we now detected rustrela virus (RusV; *Rubivirus strelense; Matonaviridae*) sequences in brains of cats with ‘staggering disease’-like neurological disorder. RusV is a recently discovered relative of rubella virus (RuV; *Rubivirus rubellae*)^32^, the causative agent of rubella in humans^30, 33^. It was first identified in the brains of various mammals in a zoo close to the Baltic Sea in Northern Germany^30, 34^. These animals had suffered from neurologic disorders and lymphohistiocytic encephalitis^30, 31, 34^. Yellow-necked field mice (*Apodemus flavicollis*) without apparent encephalitis were considered as possible reservoir hosts of the virus in that area^30, 34^.

Here, we now report the presence of RusV in the brains of cats with non-suppurative lymphohistiocytic meningoencephalomyelitis and ‘staggering disease’-like disorders from Sweden, Austria, and Germany, by metagenomic analysis and further independent methods, including reverse transcription real-time PCR (RT-qPCR), ISH and IHC. In contrast, RusV was not detected in the brains of control cats without neurologic disease or with encephalopathies of other causes from the same or nearby regions. Thus, our results indicate that RusV is indeed the causative agent of long-known ‘staggering disease’ in domestic cats.

## RESULTS

### Failure to detect BoDV-1 infection in cats with ‘staggering disease’

In an attempt to investigate the aetiology of ‘staggering disease’, frozen or formalin-fixed paraffin-embedded (FFPE) brain samples from 29 cats with non-suppurative encephalitis and/or the clinical presentation of ‘staggering disease’ from Sweden (n=15), Austria (n=9), and Germany (n=5) were examined for the presence of bornaviruses by RT-qPCR assays detecting the RNA of BoDV-1 and other orthobornaviruses^27^, (**Supplemental Table S1**), or by IHC using a monoclonal antibody targeting the BoDV-1 nucleoprotein (**Supplemental Figure S1**). Neither bornavirus RNA nor antigen could be detected.

### RusV sequences identified in cats with ‘staggering disease’ by metagenomic analysis

Selected samples were subsequently analysed using a generic metagenomic sequencing workflow^32^. In an initial analysis using blastx, sequence reads with the highest identity to RusV were identified in 14 out of 15 tested samples from these three countries (**Table 1**). Additional high throughput sequencing (HTS), assisted by target enrichment using the panRubi myBaits set v2), a newly developed set v3 achieved complete RusV genome sequences for three cats from Sweden (animals SWE_13, SWE_14 and SWE_15) and one cat from Northeastern Germany (GER_04), as well as a complete and an almost genome sequences for two cats from Austria (AUT_02 and AUT_06, respectively). The newly identified RusV sequences clearly clustered with other RusV sequences when compared to related matonaviruses (**Figure 1a**), based on amino acid (aa) sequences of the structural polyprotein (p110/sPP). The genome nucleotide (nt) sequences from Austria and Sweden formed separate phylogenetic lineages in comparison to the sequences from Germany (**Figure 1b**). While sequence GER_04 possessed at least 92.1% nt sequence identity with the previously published German RusV sequences, the minimum nt identities of the Swedish and Austrian sequences to the German sequences were only 76.7%, or 76.0%, respectively, but 80.7% to each other (**Supplemental Figure S2**). The genome organization of the newly discovered RusV sequences (**Figure 1c**) was consistent with those of previously published RusV genomes^30, 34^. Using a sliding window analysis, we identified a highly conserved region at the 5’ terminus of the RusV genomes (approximate positions 1 to 300). Regions of particularly high variability covered the intergenic region between the p200 and p110 open reading frames (ORF) as well as a stretch of the p150-encoding sequence around nt positions 2,100-2,600 (**Figure 1c**).

**Table 1.** Rustrela virus (RusV) detection in brain samples from cats with or without signs of ‘staggering disease’.

| Group | Years | n | RusV detection (positive / total animals) |  |  |  |  |
| --- | --- | --- | --- | --- | --- | --- | --- |
| Country |  |  | HTS | RT-qPCR | ISH | IHC | total <sup>a</sup> |
| Cats matching the criteria of ‘staggering disease’ |  |  |  |  |  |  |  |
| Sweden | 2017-2021 | 15 | 9/9 | 15/15 | 14/14 | 15/15 | 15/15 |
| Austria | 1991-1993 | 9 | 3/4 | 8/9 | 7/9 | 8/9 | 9/9 |
| Germany | 2017-2022 | 5 | 2/2 | 3/5 | 1/3 | 4/5 | 4/5 |
| Cats with other types of encephalitis |  |  |  |  |  |  |  |
| Germany | 2017-2020 | 8 | 0/3 | 0/8 | n.d. | 0/8 | 0/8 |
| Cats without encephalitis |  |  |  |  |  |  |  |
| Sweden | 2021-2022 | 7 | n.d. | 0/7 | n.d. | 0/1 | 0/7 |
| Austria | 2021 | 5 | n.d. | 0/5 | n.d. | n.d. | 0/5 |
| Germany | 2018-2020 | 9 | n.d. | 0/9 | n.d. | 0/9 | 0/9 |

HTS: high throughput sequencing followed by metagenomic analysis; ISH: *in-situ* hybridization using RNAscope; IHC: immunohistochemistry; n.d.: not determined
<sup>a</sup> Cats were considered RusV-positive if RusV RNA or antigen was detected by at least one of the applied methods.

**Figure 1.**
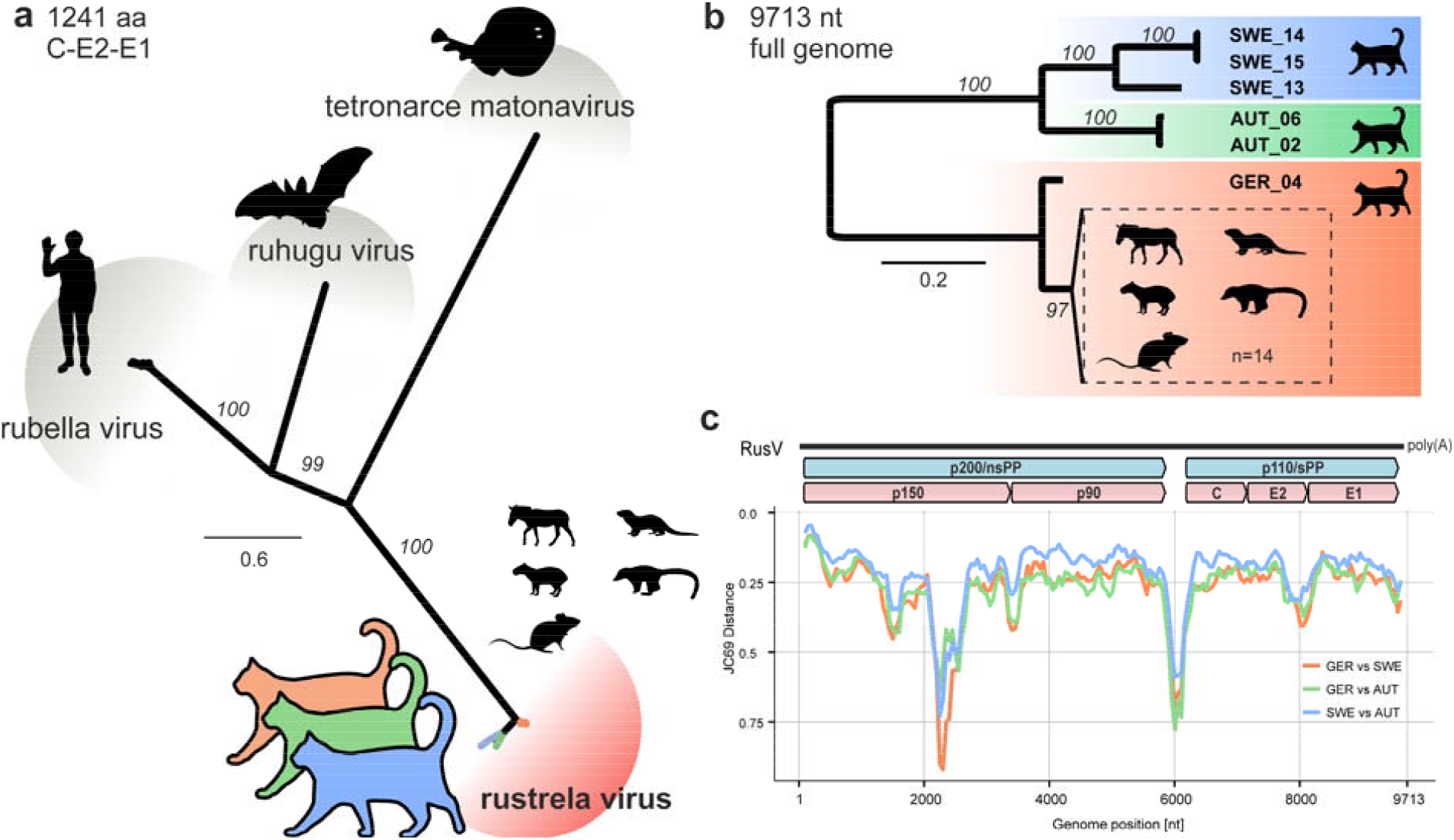
Sequence comparison of complete rustrela virus (RusV) genome sequences from cats from Sweden, Austria, and Germany. **(a)** The amino acid sequences of the structural polyprotein (p110/sPP) of all known matonaviruses were aligned and a maximum-likelihood (ML) phylogenetic tree was calculated (IQ-TREE2 version 2.2.0; FLU+F+I+G4; 100,000 ultrafast bootstraps). Bootstrap support values are shown in italics. **(b)** ML tree of complete or nearly complete RusV genome sequences from cats with ‘staggering disease’ and all publicly available RusV sequences (IQ-TREE2 version 2.2.0; TIM3+F+I; 100,000 ultrafast bootstraps). Sequences from Sweden, Austria, and Germany are highlighted in blue, green, and orange, respectively. Sequences from a previously identified German RusV cluster from zoo animals with encephalitis and apparently healthy yellow-necked field mice (*Apodemus flavicollis*)^30, 31, 34^ are presented in a dashed box. Bootstrap support values are shown at the nodes. **(c)** The genetic variability of RusV lineages from Sweden, Austria, and Germany is presented as mean pairwise JC69 distance using a sliding window analysis (window: 200 nt; step size: 50 nt). The genomic organization of RusV is shown, highlighting the non-structural (p200/nsPP) and structural (p110/sPP) polyprotein open reading frames, as well as the mature cleavage products protease (p150), RNA-directed RNA polymerase (p90), capsid protein (C), and glycoproteins E2 and E1.

### Detection of RusV RNA using a broadly reactive panRusV RT-qPCR assay

Since the initially published RT-qPCR assay RusV Mix-1^30^ was unable to detect RusV RNA in samples from Sweden and Austria (data not shown), we designed a new set of primers and probe targeting the highly conserved region at the 5’ end of the genome (**Supplemental Figure S3; Supplemental Table S1**). This newly established panRusV assay readily detected RusV RNA in the brains of all 15 Swedish cats with ‘staggering disease’, eight out of nine Austrian cats^8, 11^, and three out of five cats from Germany (**Table 1**). Results were moderately to strongly positive for frozen tissue (cycle of quantification [Cq] values 20 to 32), and rather weakly positive for animals where only FFPE material was available (Cq 27 to 36; **Supplemental Table S2; Supplemental Figure S4a**). In contrast, RusV RNA was not detected in frozen brain samples from 21 control cats without encephalitis originating from Sweden, Austria, and Germany, or in eight cats from Germany suffering from other types of encephalitis (**Table 1**).

### Detection of RusV RNA and antigen in neural tissue by ISH and IHC

To confirm and further characterize RusV infection in the cats, we employed viral RNA detection by ISH and antigen detection by IHC (**Figure 2**). An RNAscope ISH probe was designed to target the highly conserved stretch at the 5’ terminus of the RusV genome (**Supplemental Figure S3**). Specific ISH staining was observed for 22 out of 26 tested cats from all three countries (**Table 1; Figures 2a to d**). Two animals revealed inconclusive results and two were ISH-negative (**Supplemental Table S2**).

**Figure 2.**
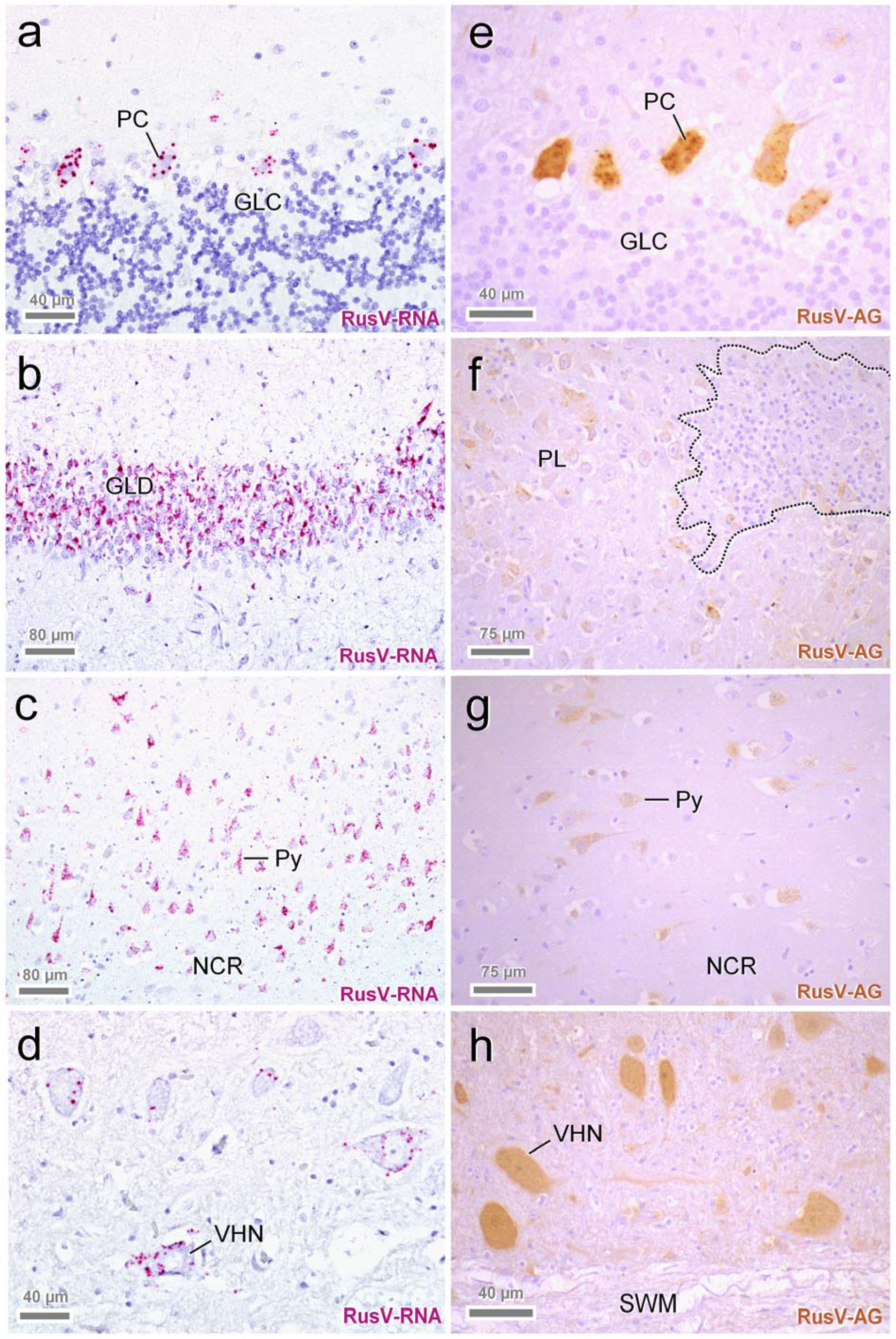
Detection of rustrela virus (RusV) RNA by RNAscope *in-situ* hybridization (a-d) and RusV antigen by immunohistochemistry (e-h) in the central nervous system of encephalitic cats. Both virus RNA and capsid protein were located mainly in the cytoplasm of different nerve cell populations. Typical are spherical reaction products, which may coalesce to more extensive and/or diffuse staining. Neurons with the highest viral load were particularly Purkinje cells (**a, e:** PC), granule cells of dentate gyrus (**b**: GLD), pyramidal cells within hippocampal pyramidal cell layer (**f**: PL), and neocortex (**c, g:** Py). Also, numerous RusV positive cells are seen in lower brainstem and spinal ventral horn neurons (**d, h**: VHN). Note that expression of virus RNA and antigen is far more widespread than inflammatory changes (**f**: dashed line) and mostly affects neurons without cytopathic effects. GLC: granule cell layer of cerebellar cortex; GLD: granule cell layer of dentate gyrus; NCR: neocortical ribbon; PC: Purkinje cell; PL: pyramidal cell layer; Py: pyramidal cell; SWM: spinal white matter; VHN: ventral horn neuron. Sources: (**a**) cat SWE_03; (**b**) SWE_11; (**c**) SWE_05; (**d, f**) AUT_09; (**e**) AUT_04; (**g**) SWE_08; (**h**) AUT_08.

In addition, we performed IHC using a newly generated mouse monoclonal antibody targeting the RusV capsid protein. Specific immunopositivity mirroring the ISH pattern (**Figures 2e to h**) was seen in 27 out of 29 analysed cats with ‘staggering disease’, but not in any brain from 18 tested control cats (**Table 1**). IHC identified RusV antigen in two cases that had been negative by RT-qPCR from FFPE brain tissue (AUT_03 and GER_02), whereas one RT-qPCR-positive individual (AUT_05) remained negative by IHC (**Supplemental Table S2**).

By both, ISH and IHC, a specific granular chromogen staining was observed predominantly in perikarya of pyramidal neurons of cerebral cortices, namely of isocortex (**Figures 2c, g**) and hippocampus proper (**Figure 2f**), granule cells of dentate gyrus (**Figure 2b**), Purkinje cells of the cerebellum (**Figures 2a, e**), multipolar neurons of brain stem and cerebellar roof, and in ventral horn neurons of the spinal cord (**Figures 2d, h**). On occasion, cytoplasmic immunoreactivity was also noted in individual interposed neuroglial and microglial cells. In addition, some small dot-like reactions were spotted in a scattered pattern amongst the neuropil and white matter. Notably, viral RNA and protein often did not colocalize with inflammatory lesions (**Figure 2f**).

### Demographic data, clinical disease and histopathology of RusV-infected cats

Among the 29 cats in this study that met the criteria of ‘staggering disease’, 28 cats were identified as RusV-positive by at least one of the employed methods (**Table 1; Supplemental Table S2**). Twenty-two (78.6%) of them were neutered or intact males (**Supplemental Table S3**), which is consistent with previous studies on ‘staggering disease’^14, 15, 35^. All affected animals were adults, with a median age of 3.1 years (range 1.5 to 12.3; **Supplemental Figure S5a; Supplemental Table S3**), and all had outdoor access (where reported) (**Supplemental Table S3**). The onset of disease had occurred more often in winter and spring (December to May: 18 cases) as compared to summer and autumn (June to November: 8 cases; **Supplemental Figure S5b; Supplemental Table S3**).

Typically observed clinical signs included gradually deteriorating gait abnormalities, with abnormal posture, stiff gait, ataxia, hind limb-predominant weakness progressing to non-ambulatory tetraparesis and proprioceptive deficits. In addition, fever, behavioural changes such as abnormal vocalization or affectionate behaviour, hypersalivation, depression, hyperaesthesia in dorsal back and lumbar/tail region, reduced spinal reflexes and postural reactions, affection of cranial nerves and opisthotonus were reported in some of the cases. In one animal, generalized seizures were specifically recorded (**Supplemental Table S4**)^6, 8^. Duration from the reported disease onset to euthanasia ranged from one week to more than one year, with most of the cats being euthanized within less than two months (median two weeks) (**Supplemental Figure S5c; Supplemental Table S3**).

In congruence with previous reports on ‘staggering disease’^8, 11, 15^, histopathological examination of brain and spinal cord revealed widespread, polio-predominant angiocentric lymphocytic and/or lymphohistiocytic infiltrates throughout the cases (**Figures 3 and 4; Supplemental Table S4**). Occasionally, they were accompanied by oligofocal astrogliosis and microglial proliferates, a few degenerating neurons and neuronophagic nodules (**Figure 4**). Inflammation was most pronounced in the brain stem (**Figures 3a, 4b, c**), cerebral cortices (**Figures 3b, c**), and all levels of the spinal cord, while – independent of the ISH and IHC signal (**Figures 2a, e**) – they were less evident in the cerebellum (Figure 3a). Apart from the parenchyma, lymphohistiocytic infiltrates and fewer plasma cells were present also in the leptomeninges (**Figure 3a**). Potentially viral inclusion bodies were not observed.

**Figure 3.**
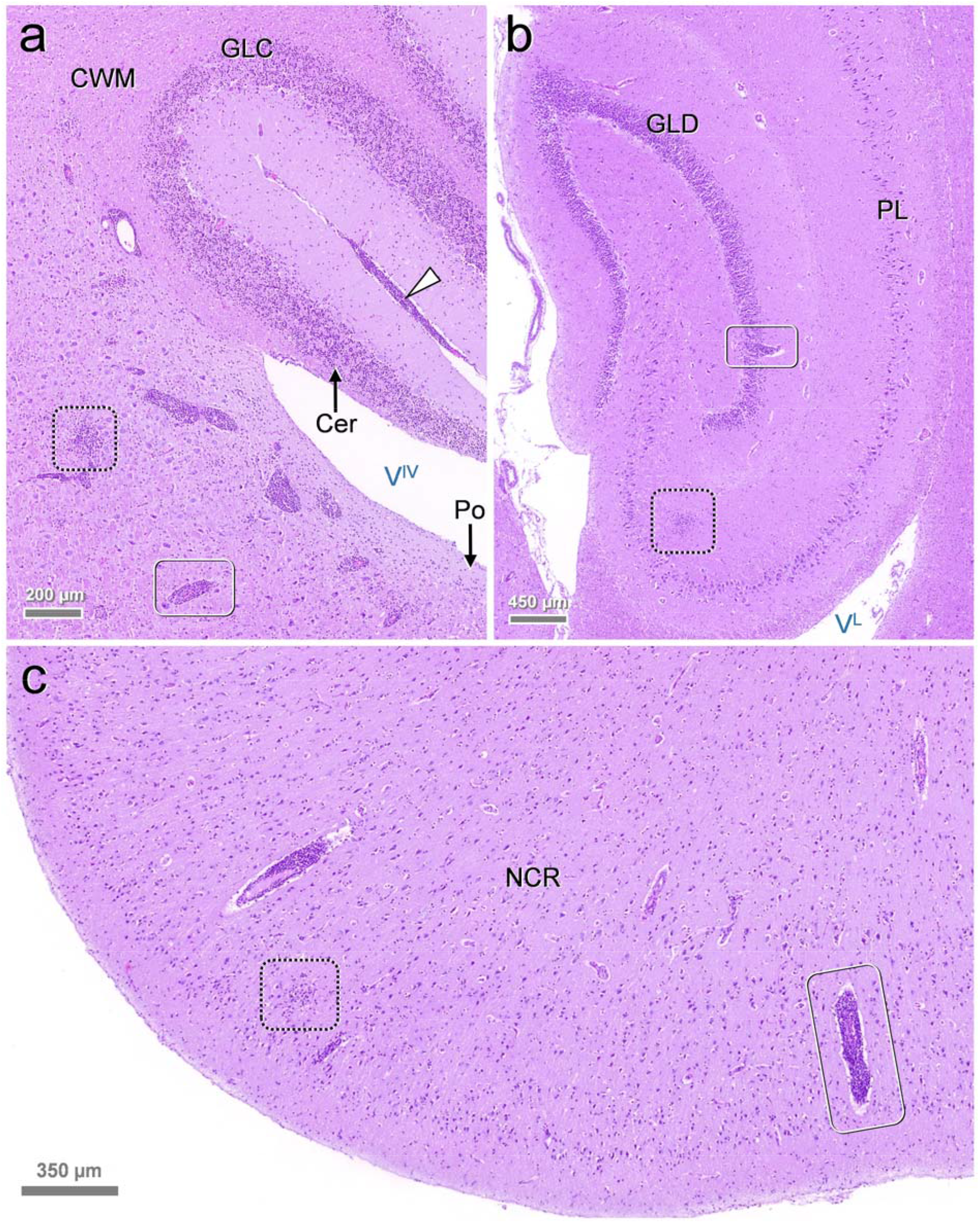
Encephalitic pattern in rustrela virus (RusV)-infected cats. Histology typically features polio-predominant, perivascular lymphohistiocytic cuffs (**a-c**: solid boxes) and angiocentric infiltrates (**a-c**: dashed boxes). They are most prominent in brainstem (**a**: Po), hippocampus formation (**b**) and neocortex (**c**). Leptomeningeal infiltrates (**a**: white arrowhead) also occur in areas with sparse parenchymal invasion such as the cerebellum (**a**: Cer). Stain: hematoxylin eosin (H.E.). Anatomical landmarks: Cer: cerebellum; CWM: cerebellar white matter; GLC: granule cell layer of cerebellar cortex; GLD: granule cell layer of dentate gyrus; NCR: neocortical ribbon; PL: pyramidal cell layer; Po: pons; V^IV^: fourth ventricle; V^L^: lateral ventricle. Sources: (**a**) cat SWE_04; (**b, c**) SWE_07.

**Figure 4.**
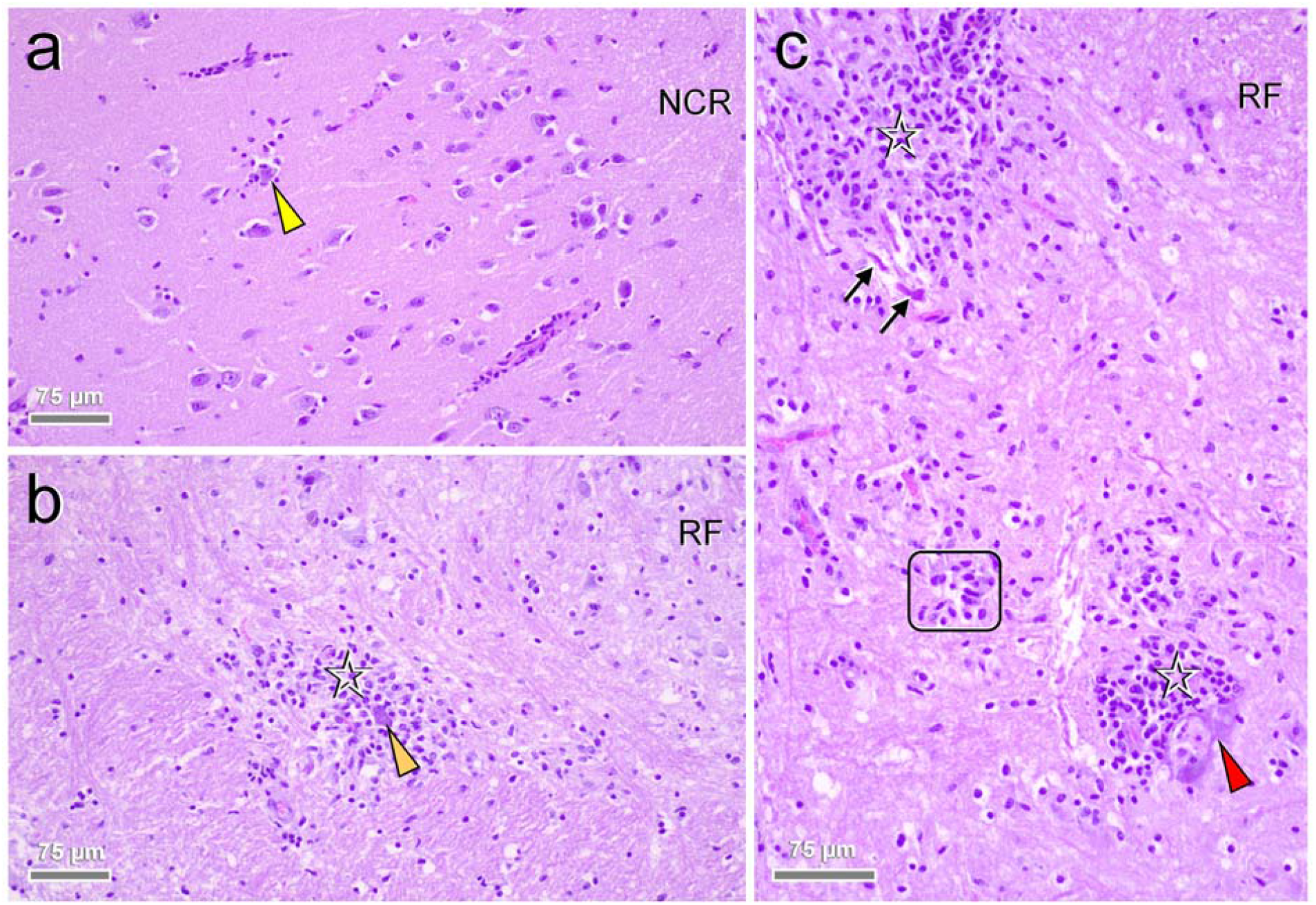
Close-up pathology and cellular damage of rustrela virus (RusV) infection within the brain. Infected brains show neurons (**a-c**: arrowheads) with (**c**: red arrowhead) and without (**a, b**: yellow and orange arrowheads) degenerative features, early (**a, b**: yellow and orange arrowheads) and advanced (**c**: red arrowhead) neuronophagia suggestive of a neuronotropic pathogen. Focal dropout of neurons goes with formation of microglial stars (**c**: frame). Inflammatory infiltrates (**b, c**: asterisks) mingle with focal glial proliferates. Dystrophic axons (**c**: black arrows) are occasionally present within the perilesional area. Stain: hematoxylin eosin (H.E.). Anatomical landmarks: NCR: neocortical ribbon; RF: reticular formation. Sources: (**a**) cat SWE_06; (**b, c**) SWE_04.

### Detection of RusV RNA in rodents from Southern Sweden

We furthermore screened brain samples from 116 rodents that had been collected between 1995 and 2019 during monitoring studies near Grimsö in Örebro county (**Supplemental Figure S6**), which is situated approximately 80 km Southwest of the origin of the closest RusV-infected cat detected in this study (**Figure 5**). PanRusV RT-qPCR detected RusV RNA in eight out of 106 (7.5%) wood mice (synonym ‘long-tailed field mice’; *Apodemus sylvaticus*) with Cq values ranging from 20 to 35 (**Supplemental Figure S4b**). In contrast, we did not detect RusV RNA in ten yellow-necked field mice from the same location. The positive individuals were collected in the years 1996 (n=2), 1997 (n=3), 2005 (n=2), and 2011 (n=1). All positive animals had been trapped during fall season, which is consistent with the considerably higher number of wood mice trapped during fall (n=94) as compared to spring (n=12; **Supplemental Figure S6**).

**Figure 5.**
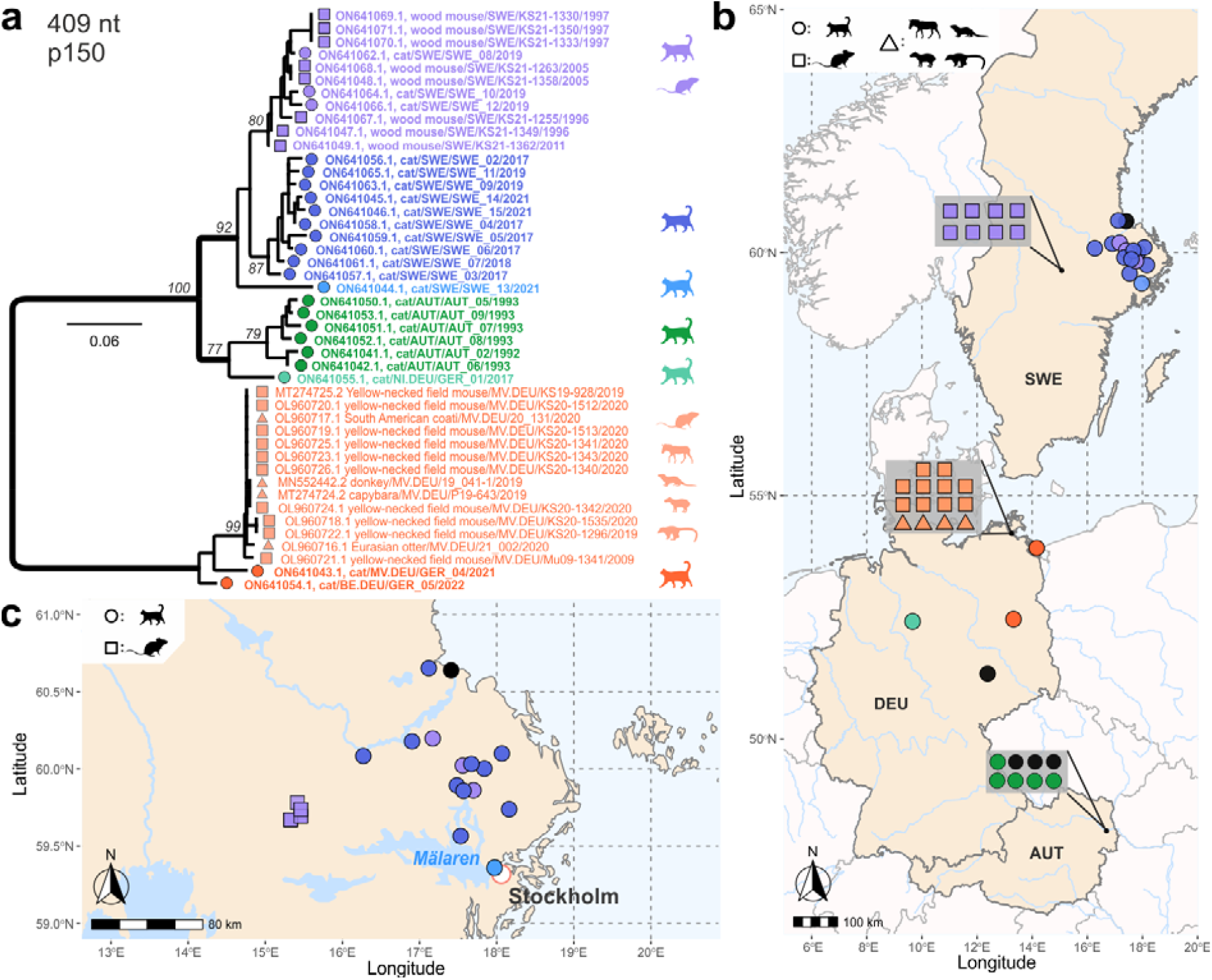
Phylogenetic analysis and spatial distribution of rustrela virus (RusV) infections in Europe. (**a**) Maximum likelihood (ML) phylogenetic tree of partial RusV sequences (409 nucleotides, representing genome positions 100 to 508 of donkey-derived RusV reference genome MN552442.2; IQ-TREE version 2.2.0; TN+F+G4; 100,000 ultrafast bootstraps). Only bootstrap values ≥70 at major branches are shown in the phylogenetic tree. RusV sequence names are shown in the format “host/ISO 1366 code of location (federal state.country)/animal ID/year”. (**b, c**) Mapping of the geographic origin of RusV-positive animals in Europe (b) and in the Lake Mälaren region in Sweden (c). Colours represent the phylogenetic clades of the sequences (a). RusV-positive cats that failed to deliver sequences are depicted in black. The respective host animals are shown as circles (cats), squares (*Apodemus* spp.), and triangles (zoo animals). Symbols in grey boxes represent individuals from the same or very close locations. AUT = Austria, DEU/GER = Germany, SWE = Sweden; BE = Berlin, MV = Mecklenburg-Western Pomerania, NI = Lower Saxony.

None of the positive animals showed inflammatory lesions in their brain tissues (data not shown). Sample quality allowed for ISH analysis of brain tissue for only four RusV-positive individuals. All of them exhibited specific staining, whereas one RT-qPCR-negative wood mouse did not when tested in parallel (**Supplemental Figure S7**).

### Phylogenetic analysis and spatial distribution of RusV sequences from cats and wood mice

To allow for a detailed phylogenetic analysis, we aimed at generating RusV sequence information for all positive cats and wood mice. However, whole RusV genome sequencing by HTS is sophisticated and laborious^34^. Particularly for those individuals with only FFPE material available, the generated sequences were highly fragmented. Thus, we designed primers specifically targeting a stretch of 409 nt within the highly conserved region at the 5’ end of the genome to be applied for conventional RT-PCR and subsequent Sanger sequencing (**Supplemental Table S1; Supplemental Figure S3**). Using this approach, sufficient sequence information was generated for 23 RusV-positive cats and all eight RusV-positive wood mice. Phylogenetic analysis of these sequences together with all previously published RusV sequences revealed three clearly distinguishable clades for sequences originating from Sweden, Austria, or Northern Germany, with the Swedish and Austrian clades being more closely related to each other than to the Northern German clade (**Figure 5a**). The Swedish clade revealed three distinguishable subclades. One subclade harboured sequences from ten cats from an area of about 9,000 km^2^ around the city Uppsala. A second subclade included three RusV sequences from cats from the same region and all sequences from wood mice from Grimsö. The third subclade was constituted by only a single sequence from a cat from Stockholm (Figure 5). The sequences of both cats from Northeastern Germany belonged to the previously published Northern German clade (**Figures 5a, b**)^30, 34^. Surprisingly, sequence fragments available for cat GER_01, which originated from Hannover in Central Germany, were more closely related to sequences of the Austrian clade than to the Northern German clade (**Figures 5a, b**).

## DISCUSSION

For almost five decades, ‘staggering disease’ in domestic cats had been suspected as a cohesive entity with a uniform, presumably viral, but still unknown aetiology^7, 8, 11, 14, 15^. While BoDV-1 had been discussed as a candidate for causing ‘staggering disease’^16, 19, 21, 22, 23^, proof of natural infections complying with current diagnostic standards could not be presented^17, 24, 25^. Here we used robust diagnostic approaches that had been demonstrated to successfully detect a broad range of orthobornaviruses, including cases of BoDV-1-induced encephalitis in humans and domestic mammals^27, 36, 37, 38^. Nevertheless, we were not able to detect bornavirus RNA in any of the 29 tested cats from three different countries with clinicopathological features consistent with ‘staggering disease’. Thus, our results clearly refute the hypothesis of BoDV-1 being the causative agent of ‘staggering disease’.

Instead, we were able to unequivocally confirm RusV infection in almost all of these cats. We consistently detected RusV RNA and antigen by employing independent diagnostic assays, including RT-qPCR, genome sequencing, ISH and IHC in the majority of these individuals. Only minor inconsistencies between results of the assays occurred, presumably due to genetic variability of the involved RusV variants, quality of the available material and differential sensitivities of the employed assays that may have led to false negative results of single tests, particularly for individuals for which only archived FFPE material was available. Experimental RusV infection of cats, to reproduce the disease and thereby fulfil Henle-Koch’s postulates, has not been performed so far due to the lack of a virus isolate. However, we demonstrate a clear association between infection and disease, with almost all animals of the ‘staggering disease’ group being RusV-positive, whereas the virus was not detected in any control cat without neurologic disease or with other types of encephalitis. Furthermore, clinical course and histopathologic lesions observed for cats with ‘staggering disease’ in this and previous studies^7, 8, 14, 15^ closely resembled those described for other RusV-infected mammals in Germany^30, 34^. Thus, RusV can be considered as the causative agent of ‘staggering disease’ with high certainty.

We detected RusV infection of cats in the Lake Mälaren region in Sweden and Northeast of Vienna in Austria, two traditional hotspots of ‘staggering disease’^7, 8, 11, 15, 35^, as well as in Northern Germany, where RusV had been initially discovered^30, 34^, but ‘staggering disease’ had not yet been reported. Phylogenetic analyses revealed the RusV sequences from the three regions to belong to separate genetic clusters, with the Swedish and Austrian sequences being more closely related to each other than to those from Northern Germany. The considerable genetic variability with only 75% minimal nt sequence identities among the different lineages posed a major challenge for the generation of broadly reactive diagnostic tools. However, a particularly conserved sequence stretch at the 5’ terminus of the genome allowed for the design and application of versatile primers and probes for PCR and ISH assays. Furthermore, a monoclonal mouse antibody targeting the RusV capsid protein proved suitable for the detection of all three major RusV lineages.

While yellow-necked field mice are considered as putative reservoir hosts of RusV in Northern Germany^30, 34^, we surprisingly detected RusV in Sweden only in the closely related wood mice but not in yellow-necked field mice from the same area in Örebro county. Since the majority of tested individuals from this location were wood mice, it remains to be elucidated, whether this discrepancy is mainly a result of different species compositions of the analysed sample collections or whether it represents a diverging adaptation of RusV variants to alternative reservoir hosts.

The route of RusV transmission within its presumed reservoir as well as from there to other hosts remains unknown. The tissue tropism in zoo animals and yellow-necked field mice in Germany was described as restricted almost exclusively to the CNS, with occasional detection of RusV RNA in peripheral nerve fibres. Viral shedding has not been described so far^30, 34^. In the future, detailed data on tissue distribution needs to be obtained also for RusV-infected cats and wood mice, but this was beyond the scope of this study. Furthermore, the possibility of RusV shedding by infected cats remains to be elucidated. However, the apparently spatially restricted occurrence of the phylogenetic clusters argues in favour of a continuous viral spread only within a locally bound reservoir, including small mammals, whereas more mobile hosts, including domestic animals that may be transported over long distances, serve predominantly as erroneous dead-end hosts. Similar patterns have been evidenced for rodent and shrew reservoir-bound viruses such as BoDV-1, rat hepatitis E virus or Puumala orthohantavirus^25, 39, 40, 41^. The sporadic occurrence of ‘staggering disease’ in domestic cat populations, the apparent lack of outbreak series within cat holdings, as well as the almost exclusive restriction to cats with outdoor access, often originating from rural areas, further support this assumption^8, 14, 15, 35^.

Previous studies had also suggested a seasonal occurrence of ‘staggering disease’ with more cases in winter and spring than summer and autumn^15^. Although higher case numbers and a more systematic sampling scheme are required for solid statistical evaluation, the same tendency was observed also in our study. This seasonal pattern may be attributable to fluctuating reservoir populations. During the small mammal monitoring in Grimsö, Sweden, numbers of *Apodemus* spp. trapped in fall were much higher than in spring. In addition, movement of small rodents towards and into human dwellings during winter is frequently reported and has been discussed to be associated with transmission of zoonotic pathogens such as Puumala orthohantavirus to humans^42^. Increased exposure to *Apodemus* spp. during fall and winter might also facilitate RusV transmission to cats. However, since the incubation period of RusV-induced disease is unknown, time points of infection cannot be reliably estimated so far. Changes of reservoir populations may also explain long-term temporal patterns of ‘staggering disease’ occurrence. While cases have been continuously observed in the Swedish Lake Mälaren region from at least the 1970s until today^7, 15, 22, 23, 35^, ‘staggering disease’ in the districts north-east of Vienna has been diagnosed mainly during the early 1990s^8, 11^, but appears to have ceased thereafter.

In summary, we provide convincing evidence of an association of RusV infection with ‘staggering disease’ in cats, strongly supporting a causative role. Our results demonstrate a much broader genetic diversity and spatial distribution of RusV than initially appreciated, and we identified the wood mouse as an additional potential reservoir host. The availability of broadly reactive diagnostic tools may lead to the detection of RusV in encephalitic cats also in regions where ‘staggering disease’ has not been evident before. Furthermore, given the broad range of affected zoo animals^30, 34^, RusV may be found to be responsible also for additional neurologic disorders in other mammalian species, possibly even including humans. Thus, future research should include the investigation of a possible zoonotic potential of RusV.

## MATERIAL AND METHODS

### Samples and data collection

Fresh-frozen or formalin-fixed paraffin-embedded (FFPE) brain and/or spinal cord samples from 29 cats fulfilling the inclusion criteria for this study (lymphohistiocytic meningoencephalomyelitis, meningoencephalitis, encephalomyelitis or encephalitis of unknown cause, and clinical signs suggestive of ‘staggering disease’) were provided by different laboratories from Sweden, Austria, and Germany (**Table 1; Supplemental Tables S2, S3, and S4**). The samples dated back to 1991 to 1993 (Austria) or 2017 to 2022 (Germany and Sweden). Some of these cases were published previously^6, 8, 11^. In addition, frozen brain samples from 21 cats originating from Sweden, Austria, and Germany without encephalitis were included as controls. An additional control group was composed of eight cats from Germany that had suffered from encephalitis of other types or causes, such as CNS manifestation of feline coronavirus (FCoV)-associated feline infectious peritonitis (FIP), vasculitic disorders and immune-mediated limbic encephalitis (**Table 1**)^6^. Metadata were provided by the submitters and/or extracted from the previous publications, including course and duration of disease, age, sex, origin, and outdoor access of the cats (**Supplemental Table S3**), as well as clinical signs (**Supplemental Table S4**).

Furthermore, the study includes archived frozen brain samples from yellow-necked field mice (*A. flavicollis*; n=10) and wood mice (*A. sylvaticus*; n=106) that had been collected near Grimsö, Örebro county, Sweden, as part of the Swedish Environmental Monitoring Program of Small Rodents at the Grimsö Wildlife Research Station^43^. Trapping was approved by the Swedish Environmental Protection Agency (latest permission: NV-412-4009-10) and the Animal Ethics Committee in Umeå (latest permissions: Dnr A 61-11), and all applicable institutional and national guidelines for the use of animals were followed. Species identities were confirmed by cytochrome b gene sequencing as described previously^44^.

### RNA extraction

Fresh-frozen samples were mechanically disrupted in 1 ml TRIzol Reagent (Life Technologies, Darmstadt, Germany) by using the TissueLyser II (Qiagen, Hilden, Germany) according to the manufacturers’ instructions. After the addition of 200 µl chloroform and a centrifugation step (14,000 × *g*, 10 min, 4°C), the aqueous phase was collected and added to 250 µl isopropanol. Total RNA was extracted using the silica bead-based NucleoMagVet kit (Macherey & Nagel, Düren, Germany) with the KingFisher™ Flex Purification System (Thermo Fisher Scientific, Waltham, MA, USA) according to the manufacturers’ instructions. *In vitro*-transcribed RNA of the enhanced green fluorescent protein (eGFP) gene was added during the extraction procedure as described by Hoffmann *et al*.^45^.

RNA extraction from FFPE tissue was performed with a combination of truXTRAC FFPE total NA Kit (Covaris, Brighton, UK) and the Agencourt RNAdvance Tissue Kit (Beckman Coulter, Krefeld, Germany). FFPE sections were loaded into a microTUBE-130 Screw-Cap (Covaris) together with 110 µl Tissue Lysis Buffer and 10 µl proteinase K solution (both Covaris). The lysate was processed with a M220 Focused ultrasonicator (Covaris) according to the manufacturer’s recommendations for acoustic pellet resuspension. The tube was subsequently incubated at 56°C in a thermal shaker at 300 rpm overnight (no longer than 18 hours). Subsequently, the sample tube was cooled to room temperature and centrifuged at 5,000 × *g* for 15 min using a microTUBE-130 centrifuge adapter. A volume of 100 µl supernatant was transferred into a clean 1.5 ml reaction tube without transferring any wax or paraffin. After another centrifugation (5 min at 20,000 × *g*), 85 µl of the lower phase with the RNA-containing tissue pellet was transferred into a clean 1.5 ml reaction tube. It was incubated at 80°C for 20 min and then cooled to room temperature, before 175 µl B1 buffer (Covaris) were added, mixed, and briefly centrifuged. Thereafter, 250 µl of 65% isopropanol were added, mixed, and briefly centrifuged. Subsequently, the preparations were further processed with the Agencourt RNAdvance Tissue Kit (Beckman Coulter) with the KingFisher™ Flex Purification System (Thermo Fisher Scientific) according to the manufacturer’s instructions.

### Metagenomic analysis and complete genome sequencing by high throughput sequencing (HTS)

Total RNA was sequenced using a universal metagenomics sequencing workflow^32^. In brief, total RNA was extracted from fresh-frozen tissue samples using a cryoPREP impactor (Covaris) along with the Agencourt RNAdvance Tissue Kit (Beckman Coulter) and a KingFisher™ Flex Purification System (Thermo Fisher Scientific). Then, 350 ng RNA per sample were reverse-transcribed into cDNA using the SuperScript IV First-Strand cDNA Synthesis System (Thermo Fisher Scientific) and the NEBNext Ultra II Non-Directional RNA Second Strand Synthesis Module (New England Biolabs, Ipswich, MA, USA). Subsequently, cDNA was processed to generate barcoded sequencing libraries as described in detail elsewhere^32^. The cDNA was fragmented to 200 base pairs (bp) length (for FFPE material) or 500 bp length (for fresh-frozen material) using an M220 Focused ultrasonicator (Covaris). Subsequent library preparation was performed as described previously, with the following modification for FFPE material during size exclusion: small fragments were retained and purified twice with 1.2× Ampure XP Beads (Beckman Coulter). Libraries were quantified with the QIAseq Library Quant Assay Kit (Qiagen) and sequenced on an Ion Torrent S5XL instrument using Ion 530 chips and chemistry for 400 bp reads, or Ion 540 chips and chemistry for 200 bp reads (Thermo Fisher Scientific) for fresh-frozen or FFPE material, respectively. In addition to the original sequencing libraries, 7 µl of the libraries were used to apply a capture enrichment with the panRubi v2 myBaits panel as described elsewhere^34^. For samples with expected major sequence divergence (>20%) from the initially available RusV sequences from Northern Germany that were used for designing the panRubi v2 myBaits panel, a hybridization temperature of 61°C was used for 24-26 hours. In addition, a new panRubi myBaits panel was designed (v3) adding preliminary genome information from samples of Sweden and Austria to the v2 panel. The panRubi v3 myBaits panel consists of 19,982 baits (60-nt oligonucleotides arranged every 20 nt, 3x tiling; GC content of 67.3%) and was collapsed at 98% sequence identity. This panel was applied with a hybridization temperature of 64°C.

For selected RusV-positive samples, we additionally applied a depletion protocol in order to decrease the amount of host-derived ribosomal RNA (rRNA) within the total RNA and thereby increase the virus-to-background ratio. In detail, we used a Pan-Mammal riboPOOL reaction kit (siTOOLs Biotech, Planegg, Germany) for 0.2 and 1 µg total RNA following the manufacturer’s instructions. The rRNA-depleted RNA was then used for strand-specific library preparation with the Collibri Stranded RNA Library Prep Kit (Thermo Fisher Scientific). The libraries were checked for sufficient quality and quantity using the 4150 TapeStation System (Agilent Technologies, Santa Clara, CA, USA) with the High Sensitivity D1000 ScreenTape and reagents (Agilent Technologies) as well as a Qubit Fluorometer (Thermo Fisher Scientific) along with the dsDNA HS Assay Kit (Thermo Fisher Scientific). Pooled libraries were sequenced using a NovaSeq 6000 (Illumina, San Diego, CA, USA) running in 100 bp mode.

### *De novo* assembly and sequence annotation of HTS-derived sequences

The raw sequences from Ion Torrent and Illumina systems were processed as described previously^34^. Briefly, the platform-specific adapters were initially removed from the reads and the sequences were trimmed according to their quality using either 454 Sequencing Systems Software (version 3.0) or Trim Galore (version 0.6.6)^46^ with automated adapter selection, for Ion Torrent and Illumina reads, respectively. Subsequently, the reads were filtered according to their average G+C content using PRINSEQ-lite (version 0.20.4)^47^ with a G+C threshold of ≥60 mol%. The trimmed and filtered reads were used for *de novo* assembly using SPAdes genome assembler (version 3.15.2)^48^ running in single cell mode (--sc) and Ion Torrent mode (--iontorrent) as required. Subsequently, all contigs were mapped back to an appropriate RusV reference sequence using Geneious generic mapper with medium sensitivity (Geneious Prime 2021.0.1; Biomatters, Auckland, New Zealand), and a consensus sequence was generated. The final sequence was annotated according to an appropriate RusV reference genome using ORF detection as provided by Geneious Prime 2021.0.1.

### Bornavirus and RusV RT-qPCRs and design of adapted broad range RusV-specific primers and probes

Two RT-qPCR assays were applied for the detection of either a broad range of orthobornaviruses (panBorna v7.2; **Supplemental Table S1**) or specifically BoDV-1 (BoDV-1 Mix-1; **Supplemental Table S1**) following previously published procedures^27, 38^. Initial screening for RusV-specific RNA was performed using a TaqMan-based RT-qPCR assay (RusV Mix-1; Supplemental Table S1) targeting the initially discovered RusV sequences from a zoo in Northern Germany as described by Bennett *et al*.^30^. The exogenously supplemented eGFP RNA was amplified as RNA extraction control as described previously^45^.

To establish a new RT-qPCR assay for the detection of a broader range of RusV sequences, all available sequences from Northern Germany and Sweden were aligned and a set of primers and probe (panRusV-2; **Supplemental Table S1; Supplemental Figure S3**) was designed to target a highly conserved region at the 5’ terminus of the genome. RT-qPCR was performed with AgPath-ID One-Step RT-PCR reagents (Thermo Fisher Scientific), panRusV-2 primers (final concentration: 0.8 µM each) and probe (0.4 µM), eGFP primers (0.2 µM) and probe (0.15 µM), and 2.5 µl extracted RNA in a total volume of 12.5 µl. The reaction was performed with the following cycler setup: 45°C for 10 min, 95°C for 10 min, 45 cycles of 95°C for 15 sec, 60°C for 30 sec and 72°C for 30 sec. A standard preparation of a RusV-positive donkey brain^30^ served as positive control and was used for the calibration of Cq values in each RT-qPCR analysis.

### Determination of partial p150-encoding RusV sequences by Sanger sequencing

Highly conserved primer binding sites in the same alignment as described above were also identified for the amplification of 449 nt at the 5’ end of the p150-encoding sequence by conventional RT-PCR (**Supplemental Table S1; Supplemental Figure S3**). RNA extracted from frozen brain samples from all cats and rodents with positive panRusV RT-qPCR results was analysed using the following One-Step RT-PCR conditions: 2.5 μl RNA were amplified in a total volume of 25 μl using the SuperScript III One-Step RT-PCR system with Platinum Taq DNA polymerase (Thermo Fisher Scientific) and 0.4 µM each of primers RusV_80+ and RusV_528-(**Supplemental Table S1; Supplemental Figure S3**). The cycler setup consisted of 50°C for 30 min, 94°C for 2 min, followed by 40 cycles of 94°C for 30 sec, 63°C for 30 sec, and 68°C for 25 sec, and a final elongation step at 68°C for 5 min. Following separation and visualization by gel electrophoresis, amplification products were purified using Zymoclean Gel DNA Recovery Kit (Zymo Research, Freiburg, Germany) and Sanger sequencing service was provided by Microsynth Seqlab (Balgach, Switzerland). Amplicons were sequenced in both directions and consensus sequences of 409 bp lengths were generated after *de novo* assembly of quality- and primer-trimmed raw sequences in Geneious Prime 2021.0.1.

### Phylogeny and geographic mappings

Phylogenetic analysis of RusV sequences generated in this study was performed together with representative sequences of all currently known matonaviruses^30, 49^, as well as all publicly available RusV sequences from the INSDC database^30, 34^. For the phylogeny within the known matonaviruses, the aa sequences of the sPP were aligned using MUSCLE (version 3.8.425)^50^ with a maximum of 100 iterations. A maximum likelihood (ML) phylogenetic tree was then calculated using IQ-TREE2 (version 2.2.0)^51^ running in automatic model selection mode (FLU+F+I+G4) and applying 100,000 ultra-fast bootstrap replicates^52^. For phylogenetic analysis of RusV nt sequences, the complete or nearly complete RusV sequences were aligned using MAFFT (version 7.450)^53^. A ML tree was then calculated as described above (model: TIM3+F+I). The alignment was further used for sequence comparison with a sliding window approach that calculated the pairwise distances (Jukes Cantor 1969 model) within a window of 200 nt every 50 nt. A phylogenetic tree of partial p150 protein-coding sequences of 409 nt length was built as described above (model: TN+F+G4).

### Histopathological examination

Brain and spinal cord samples of cats were harvested on *post mortem* examination via extensive craniectomy-laminectomy. For histology, brain and spinal cord tissues from cats as well as brain tissue from all eight RusV-positive wood mice were fixed in 10% neutral-buffered formalin. Fixed neural tissues were routinely sampled, processed in an automatic tissue processor, embedded in paraffin wax, sectioned at 2–4 μm, and stained with histological standard stains including haematoxylin-eosin (H.E.), Nissl and Luxol-Fast-blue stains.

Slices were microscopically examined for the presence of non-suppurative, lymphohistiocytic encephalitis, meningoencephalitis and/or meningoencephalomyelitis. Inflammation was graded mild to severe based on the extent of inflammatory cell infiltrates. Mild encephalitis comprised few perivascular infiltrates, most of which showed one to two layers of cells and were not necessarily present in all investigated locations. One or two larger infiltrates in a single location were allowed to occur in this category. Moderate encephalitis comprised several infiltrates per location, showing three to five layers of cells, allowing single locations with larger or smaller infiltrates. Severe encephalitis comprised many perivascular infiltrates, most of which showed several layers of cells (>5) in the majority of investigated locations.

### Detection of RusV-specific RNA by *in-situ* hybridization (ISH)

A custom-designed RNAscope probe was provided by Advanced Cell Diagnostics (Newark, NJ, USA) based on the consensus sequence of the available RusV sequences from Sweden, targeting the highly conserved region at the 5’ end of the RusV genome. A probe targeting the messenger RNA (mRNA) of the ubiquitous, widely expressed housekeeping gene peptidyl-prolyl-isomerase-B (*Felis catus*-PPIB; cat. no. 455011) was used as positive control, while a probe targeting bacterial dihydropicolinate reductase (DapB; cat. no. 310043) was used as a negative control probe. Viral nucleic acid was determined using ISH with the RNAscope® 2.5 High Definition RED assay (Advanced Cell Diagnostics, Hayward, CA, USA) according to the manufacturer’s instructions. Briefly, brain slides were deparaffinized and pre-treated with 1× Target Retrieval solution and RNAscope® Protease Plus solution prior to hybridization with the target probe. Subsequently, the tissue was treated with a series of pre-amplifiers and amplifiers followed by the application of a chromogenic substrate. The samples were counterstained with Hematoxylin Gill No. 2 (Merck, Darmstadt, Germany).

Brain sections of a RusV-positive capybara^30^ served as positive control and showed positive reactivity with the specific RusV RNAscope probe. A brain sample from a RusV-negative control cat incubated with the RusV RNAscope probe and a brain sample from a RusV-positive cat incubated with an irrelevant RNAscope probe (*Mycoplasma hyopneumoniae*) served as negative controls and yielded no reactivity (data not shown). The scoring of the signals was performed as described in **Supplemental Table S5**.

### Detection of RusV and BoDV-1 antigen by immunohistochemistry (IHC) staining

Brain slides were evaluated for expression of RusV capsid protein using a mouse monoclonal primary antibody (2H11B1; see below). The slides were deparaffinised and underwent antigen retrieval in the microwave (750 W, 20 min) being immersed in 10 mM citrate buffer (pH 6.0) before incubation with the primary antibody (dilution 1:100) at 4°C for 18 hours. Successful labelling was demonstrated using ImmPRESS® polymer anti-mouse IgG (LINARIS Biologische Produkte, Dossenheim, Germany) coupled to peroxidase and a diaminobenzidine tetrahydrochloride staining kit (ImmPACT DAB substrate HRP; BIOZOL Diagnostica, Eching, Germany) according to the manufacturers’ instructions. After peroxidase reaction, sections were counterstained with hematoxylin. Sections of ae RusV-positive capybara brain^30^ served as virus-positive tissue control, whereas brain sections of cats from the control groups that had been tested negative for RusV by RT-qPCR served as negative tissue control (**Supplemental Figure S8**). Specificity of the anti-mouse IgG polymere was evaluated by two sections each of capybara brain and of PCR-confirmed RusV positive cat SWE_07, in which 2H11B1 antibody was replaced by horse serum and by anti-FCoV mouse monoclonal (FIPV 3-70, LINARIS Biologische Produkte, Dossenheim, Germany; **Supplemental Figure S8**). The scoring of the signals was performed as described in **Supplemental Table S5**.

Cat brain slides were furthermore assessed for the expression of BoDV-1 nucleoprotein using murine monoclonal antibody Bo18^54^ with the ABC detection kit (biotinylated goat anti-mouse IgG; BIOZOL Diagnostica) and diaminobenzidine tetrahydrochloride (ImmPACT DAB substrate HRP; BIOZOL Diagnostica). Brain slides of a horse confirmed as BoDV-1-infected by RT-qPCR served as positive control. Replacement of Bo18 antibody by an irrelevant mouse monoclonal antibody (FIPV 3-70) was used as negative reagent control on BoDV-1-positive horse tissue and RusV-positive feline brain SWE_07.

### Recombinant protein production and generation of a monoclonal anti-RusV capsid protein antibody

A synthetic DNA string fragment encoding aa 128 to 308 of the RusV capsid protein, based on the sequence from an infected donkey from Northern Germany (accession number MN552442.2), was ordered from GeneArt synthesis (Thermo Fisher Scientific) and inserted into the pEXPR103 expression vector (IBA Lifesciences, Göttingen, Germany) in-frame with a Strep-tag-coding sequence at the 3’end. The protein with a C-terminal Strep-tag was expressed in Expi293 cells (Thermo Fisher Scientific) and subsequently purified using Strep-Tactin XT Superflow high capacity resin (IBA Lifesciences) following the manufacturer’s instructions.

For monoclonal antibody generation, two female BALB/c mice were immunized intraperitoneally with 20 μg of purified capsid protein in compliance with the national and European legislation, with approval by the competent authority of the Federal State of Mecklenburg-Western Pomerania, Germany (reference number: 7221.3-2-042/17). The immunization, as well as the generation of hybridoma cells were performed as described previously^55^.

## ACKNOWLEDGEMENTS

The authors like to thank Arnt Ebinger, Jessica Geers, Viola Haring, Weda Hoffmann, Jenny Lorke, Ole Frithjof Pietsch, Patrick Slowikowski, Kathrin Steffen, Patrick Zitzow (all Greifswald– Insel Riems, Germany), Lisa Pichl, Sandra Aumiller (both Munich), Albin Norman and Vidar Skullerud (both Uppsala, Sweden) for their excellent technical assistance, Carola Wolf (Rostock, Germany), Josef Zoher (Deutsch-Wagram, Austria), Jonas Johansson Wensman and Ann-Catrin Hagblom (both Uppsala, Sweden) for sample submission, and Angele Breithaupt (Greifswald – Insel Riems, Germany) for providing advice and positive controls for *in-situ* hybridization and immunohistochemistry.

Part of the work was funded by the German Federal Ministry of Education and Research (BMBF), project RubiZoo (grant no. 01KI2111), donated to D.R. The work of C.W. was supported by the German Federal Ministry of Food and Agriculture through the Federal Office for Agriculture and Food (BMEL), project ZooSeq, (grant no. 2819114019). The contribution by F.E. and B.H. was funded by the Swedish Environmental Protection Agency through the Swedish Wildlife Management Fund (grant no. 2020-00093).

## DATA AVAILABILITY

The novel RusV sequences have been made publicly available in the INSDC database under the accession numbers ON641041 to ON641071.

## AUTHOR CONTRIBUTIONS

K.M., M.B., C.L., and D.R. initiated the study. K.M., H.W., S.T., and C.L. evaluated the histopathology. F.P., C.W., and B.L. performed metagenomics sequencing. J.K., P.S., P.We. and D.R, established and performed PCR assays and Sanger sequencing. F.P. and D.R. performed phylogenetic analyses. H.W., C.W.L., and J.M. established and performed RNAscope *in-situ* hybridization. K.M. and M.R. established and performed immunohistochemistry. A.A. and S.R. generated the RusV-specific monoclonal antibody. K.M., H.W., S.T., F.E., S.N., J.N., P.Wo., L.M., C.B., K.M.O., C.R., T.F., B.H., and C.L. provided samples from cats or rodents together with clinical and pathological diagnosis and metadata. K.M., F.P., H.W., C.W., and C.L. wrote parts of the manuscript. D.H., R.G.U, N.N., and M.B. supervised parts of the study. D.R. conceptualized the study and wrote and finalized the manuscript. The manuscript was critically reviewed through the contributions of all authors.

## COMPETING INTERESTS STATEMENT

The authors declare to have no competing interests.

## Notes

### Competing Interest Statement

The authors have declared no competing interest.

