## Supplemental Material for "Mystery of fatal ‘Staggering disease’ unravelled: Novel rustrela virus causes severe encephalomyelitis in domestic cats"

Table of contents:

Supplementary Figures S1 to S8

Supplementary Tables S1 to S5

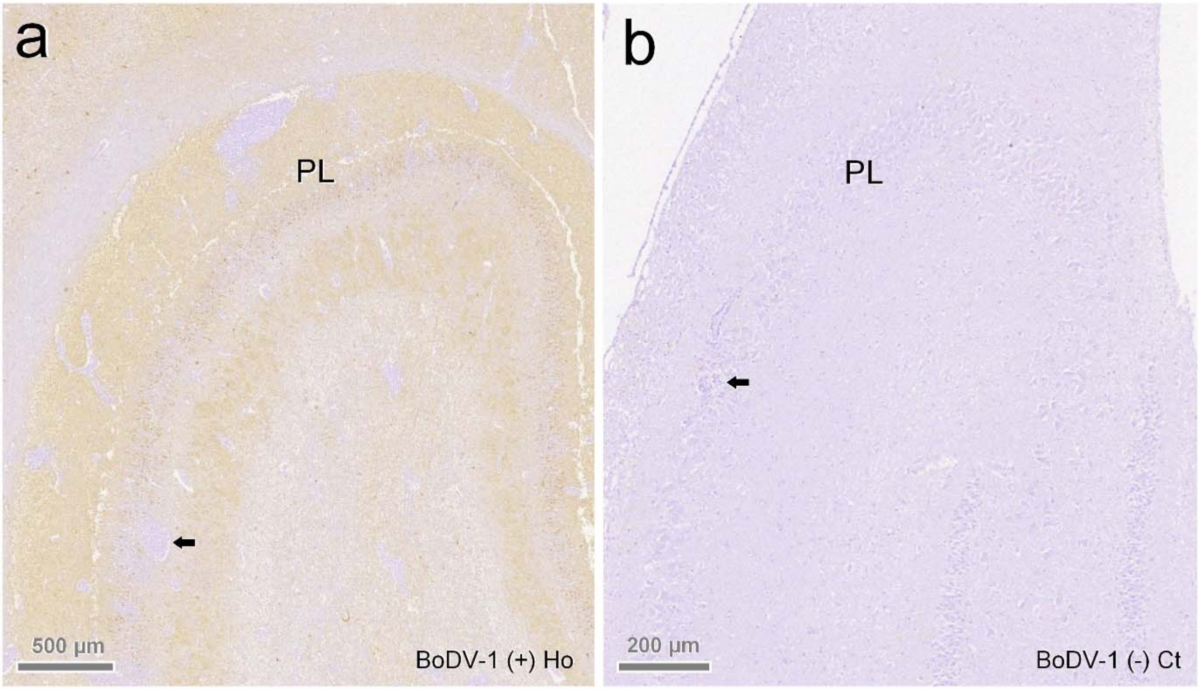


**Supplemental Figure S1. Absence of Borna disease virus 1 (BoDV-1) antigen in brains of cats with 'staggering disease'.** (**a**) Horses with Borna disease encephalitis show extensive immunopositivity for BoDV-1 nucleoprotein particularly in the hippocampus. (**b**) No BoDV-1 staining was seen in any of the tested cats, even with inflammatory infiltrates extending into hippocampus (arrow).

Ho: horse; Ct: cat; PL: pyramidal cell layer of hippocampus; arrows: angiocentric infiltrates. Source: (**b**) cat SWE_01.

| | | | 1<br>MN552442.2 | 2<br>MT274724.2 | 3<br>OL960721.1 | 4<br>OL960716.1 | 5<br>OL960722.1 | 6<br>**GER_04** | 7<br>**AUT_02** | 8<br>**AUT_06** | 9<br>**SWE_13** | 10<br>**SWE_14** | 11<br>**SWE_15** | 12<br>**KS21-1349** | 13<br>**KS21-1362** | 14<br>**KS21-1358** |
|---|---|---|---|---|---|---|---|---|---|---|---|---|---|---|---|---|
| 1 | MN552442.2 | donkey/MV.DEU(19_041-1/2019 | | 99.80 | 99.30 | 99.00 | 97.70 | 92.10 | 76.60 | 75.80 | 76.80 | 76.80 | 76.70 | 77.00 | 77.10 | 76.90 |
| 2 | MT274724.2 | Capybara/MV.DEU/P19-643/2019/Germany | 99.80 | | 99.20 | 98.90 | 97.70 | 92.10 | 76.50 | 75.70 | 76.80 | 76.70 | 76.70 | 77.00 | 77.00 | 76.80 |
| 3 | OL960721.1 | yellow-necked field mouse/MV.DEU/Mu09-1341/2009 | 99.30 | 99.20 | | 99.00 | 97.70 | 92.10 | 76.60 | 75.90 | 76.80 | 76.80 | 76.80 | 77.00 | 77.10 | 76.90 |
| 4 | OL960716.1 | Eurasian otter/MV.DEU/21_002/2020 | 99.00 | 98.90 | 99.00 | | 97.40 | 92.00 | 76.70 | 75.80 | 76.80 | 76.70 | 76.60 | 77.00 | 77.10 | 76.80 |
| 5 | OL960722.1 | yellow-necked field mouse/MV.DEU/KS20-1296/2020 | 97.70 | 97.70 | 97.70 | 97.40 | | 92.20 | 76.50 | 75.70 | 76.80 | 76.80 | 76.70 | 76.90 | 76.90 | 76.70 |
| 6 | **ON641043** | **cat/MV.DEU/GER_04/2021** | 92.10 | 92.10 | 92.10 | 92.00 | 92.20 | | 76.40 | 75.70 | 77.00 | 76.50 | 76.40 | 76.80 | 76.80 | 76.80 |
| 7 | **ON641041** | **cat/AUT/AUT_02/1992** | 76.60 | 76.50 | 76.60 | 76.70 | 76.50 | 76.40 | | 97.50 | 82.40 | 81.60 | 81.70 | 82.00 | 82.10 | 82.10 |
| 8 | **ON641042** | **cat/AUT/AUT_06/1993** | 75.80 | 75.70 | 75.90 | 75.80 | 75.70 | 75.70 | 97.50 | | 81.70 | 80.70 | 80.70 | 81.20 | 81.20 | 81.20 |
| 9 | **ON641044** | **cat/SWE/SWE_13/2021** | 76.80 | 76.80 | 76.80 | 76.80 | 76.80 | 77.00 | 82.40 | 81.70 | | 85.90 | 85.90 | 86.10 | 86.00 | 86.10 |
| 10 | **ON641045** | **cat/SWE/SWE_14/2021** | 76.80 | 76.70 | 76.80 | 76.70 | 76.80 | 76.50 | 81.60 | 80.70 | 85.90 | | 99.50 | 86.50 | 86.50 | 86.60 |
| 11 | **ON641046** | **cat/SWE/SWE_15/2021** | 76.70 | 76.70 | 76.80 | 76.60 | 76.70 | 76.40 | 81.70 | 80.70 | 85.90 | 99.50 | | 86.70 | 86.60 | 86.70 |
| 12 | **ON641047** | **wood mouse/SWE/KS21-1349/1996** | 77.00 | 77.00 | 77.00 | 77.00 | 76.90 | 76.80 | 82.00 | 81.20 | 86.10 | 86.50 | 86.70 | | 99.10 | 97.00 |
| 13 | **ON641048** | **wood mouse/SWE/KS21-1362/2011** | 77.10 | 77.00 | 77.10 | 77.10 | 76.90 | 76.80 | 82.10 | 81.20 | 86.00 | 86.50 | 86.60 | 99.10 | | 96.70 |
| 14 | **ON641049** | **wood mouse/SWE/KS21-1358/2005** | 76.90 | 76.80 | 76.90 | 76.80 | 76.70 | 76.80 | 82.10 | 81.20 | 86.10 | 86.60 | 86.70 | 97.00 | 96.70 | |

**Supplemental Figure S2. Nucleotide sequence identity matrix of an alignment of complete or nearly complete rustrela virus (RusV) genome sequences including five sequences established in this study.** Sequences generated during this study are depicted in bold. Due to the high genetic similarity of the previously published sequences from Northeastern Germany[1, 2, 3], only six out of 14 available sequences are shown. RusV sequence names are shown in the format "host/ISO 1366 code of location (federal state.country)/animal ID/year".

AUT: Austria; DEU/GER: Germany; SWE: Sweden; MV: Mecklenburg-Western Pomerania.

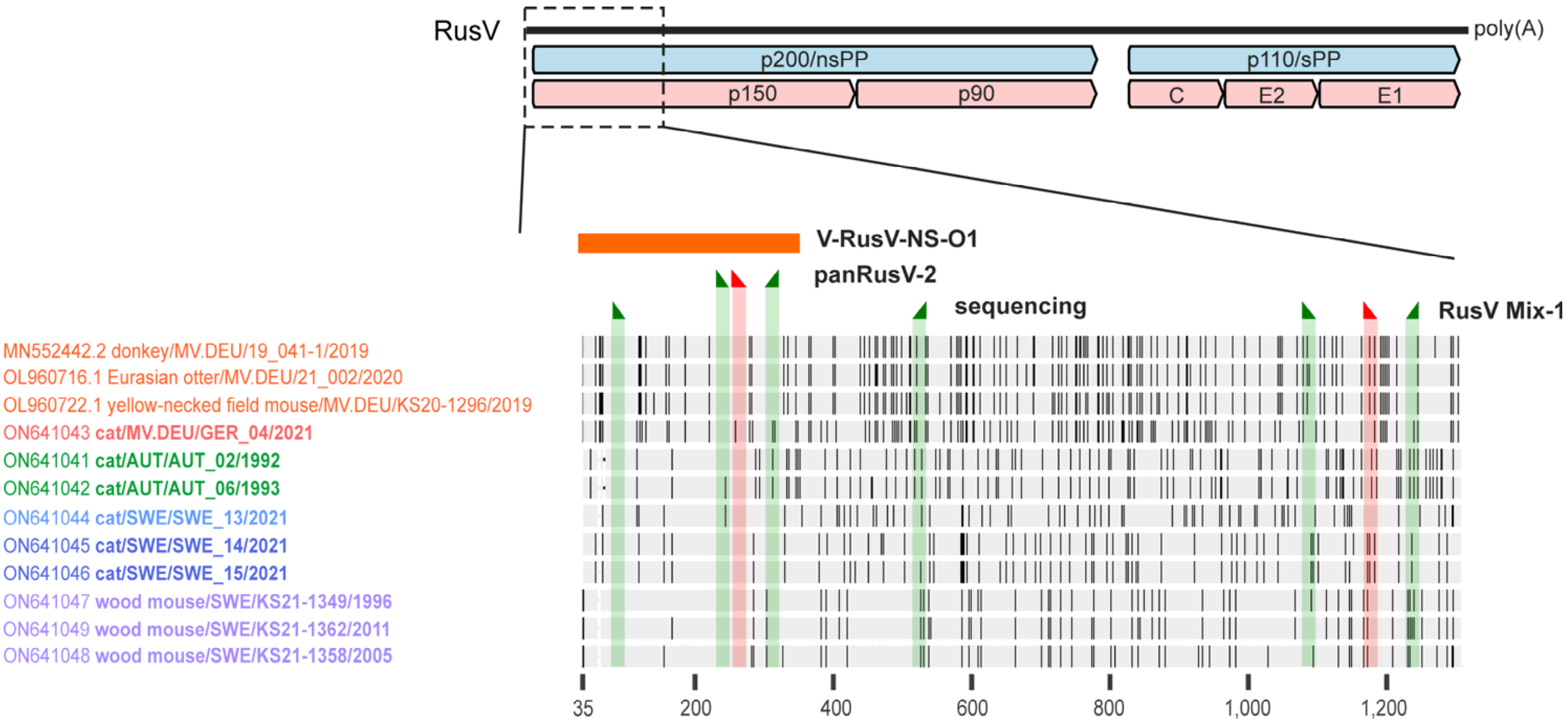


**Supplemental Figure S3. Position of primers and probes used for rustrela virus (RusV) RNA detection and sequencing.** An alignment of representative RusV sequences from Germany, Sweden, and Austria was used to visualize nucleotide mismatches in each strain (black bars) to the consensus sequence. Only the 5' end, comprising the 5' UTR and parts of the p200 non-structural polyprotein (nsPP) encoding sequence of the RusV genomes, is shown. The numbering refers to the respective genome positions in the alignment. The positions of the primers and probes of previously published RT-qPCR assays RusV Mix-1[1], and panRusV-2 designed in this study, as well as those primers used for Sanger sequencing are highlighted as green (primers) and red (TaqMan probes) triangles. The orange box indicates the stretch of the consensus of the Swedish sequence type that was used to design RNAscope probe V-RusV-NS-O1.

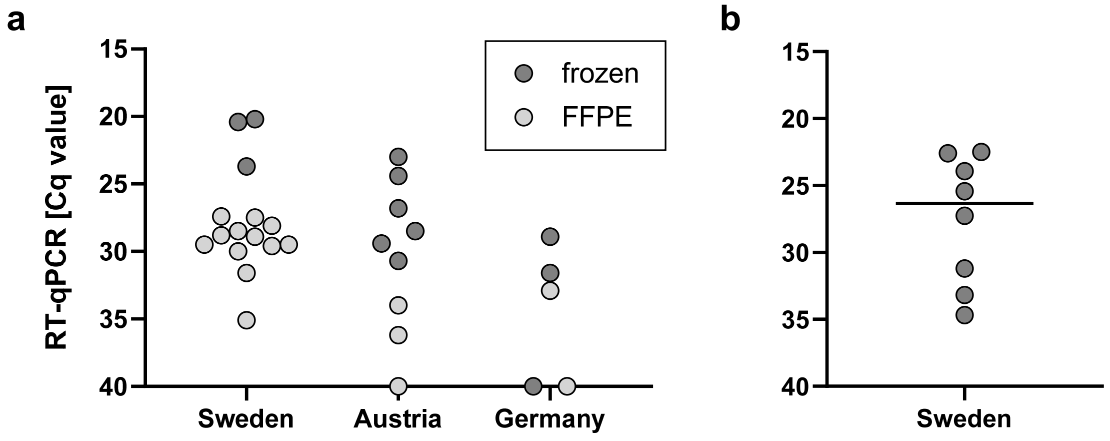


**Supplemental Figure S4. Detection of rustrela virus (RusV) RNA by panRusV RT-qPCR in brain samples.** (**a**) Cats meeting the inclusion criteria for 'staggering disease'. (**b**) Free-ranging wood mice (*Apodemus sylvaticus*) collected during monitoring studies in Grimsö (Örebro county) in Sweden. Only animals with positive results are depicted for this panel.

Cq: cycle of quantification; FFPE: formalin-fixed paraffin-embedded tissue.

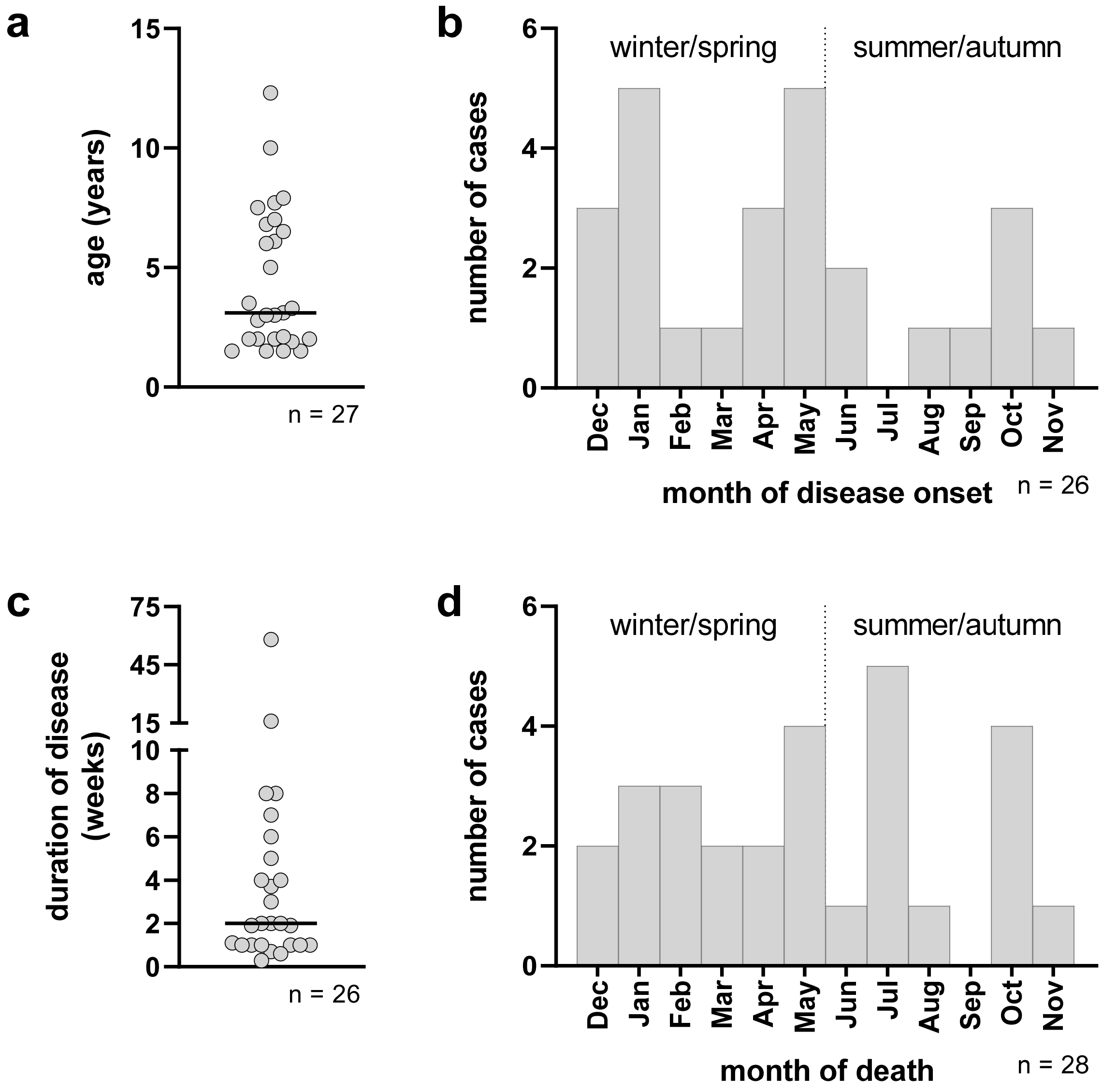


**Supplemental Figure S5. Demographic data of 28 rustrela virus (RusV)-infected cats from Sweden, Austria, and Germany.** (**a**) Age and (**c**) duration of disease as reported by the submitting veterinarian. (**b, d**) Seasonal distribution of disease onset and month of death of RusV-infected cats. The age, month of disease onset, and duration of disease were unknown for one, two, and two cats, respectively.

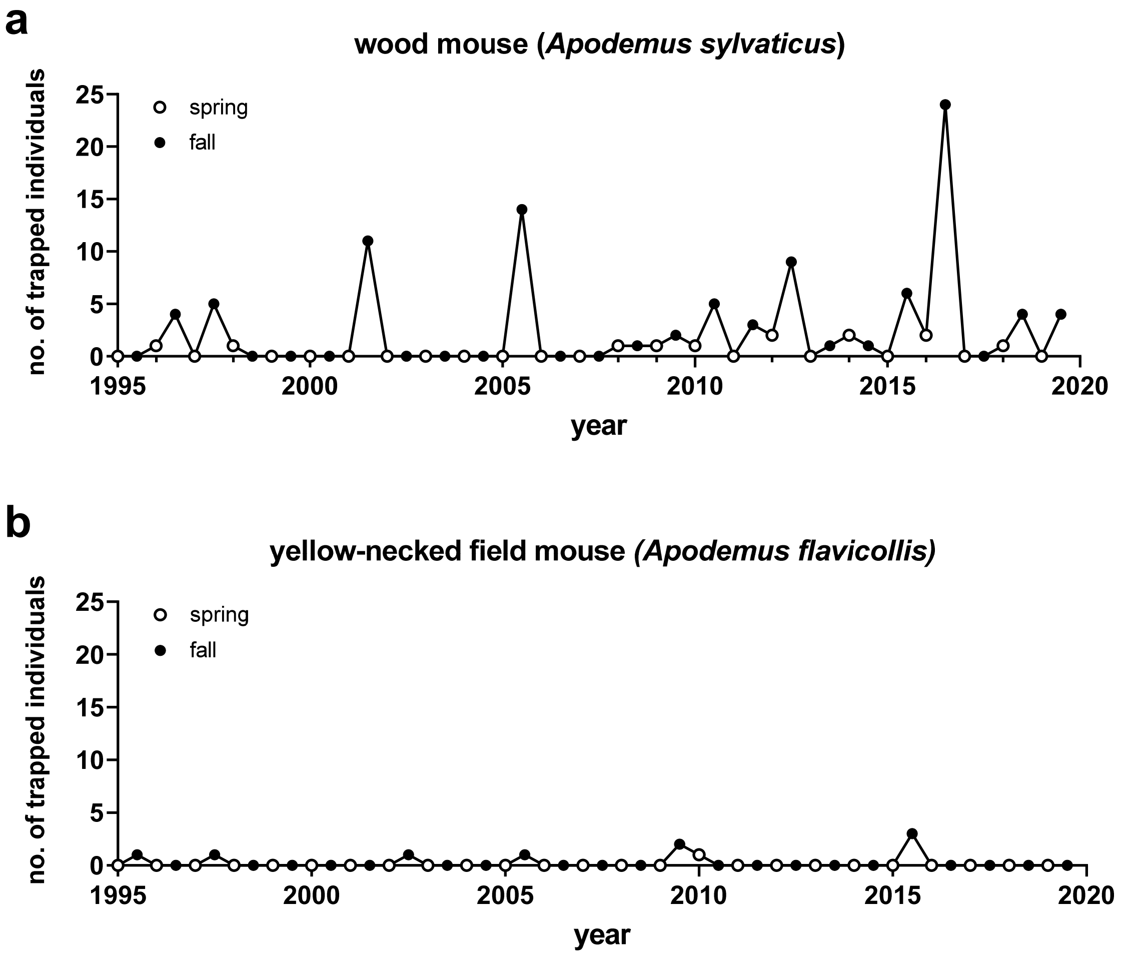


**Supplemental Figure S6. Number of trapped rodents (*Apodemus* spp.) from Grimsö, Sweden, examined during this study.** The animals had been collected as part of the Swedish Environmental Monitoring Program of Small Rodents[4] during 2,610 to 2,895 trap nights per year and season.

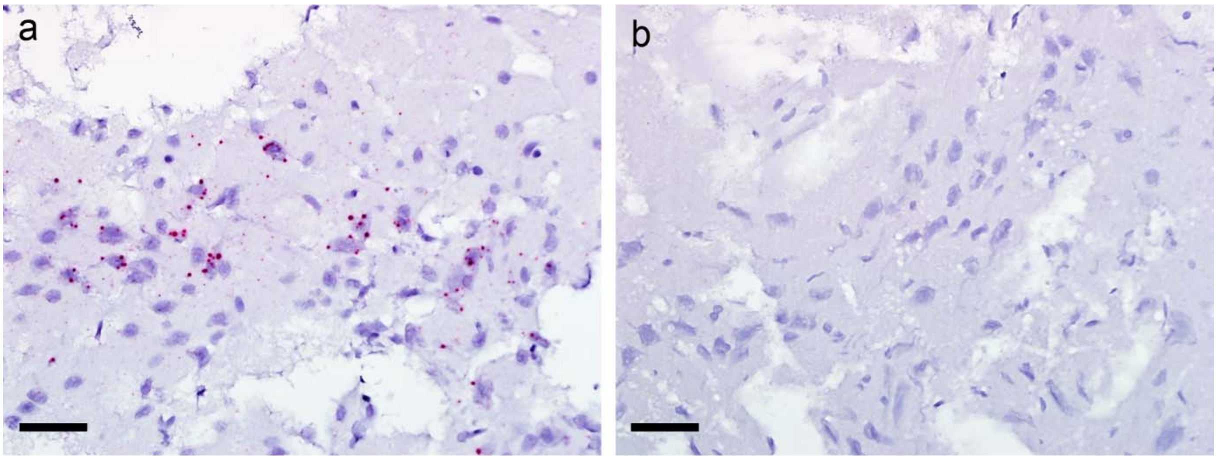


**Supplemental Figure S7. Localization of rustrela virus (RusV) RNA by RNAscope *in-situ* hybridization in the brain of wood mice (*Apodemus sylvaticus*).** (**a**) Abundant, predominately spherical reaction products in neurons and neuropil of the cerebral cortex of an RT-qPCR-positive wood mouse. (**b**) No reactivity in an RT-qPCR-negative wood mouse. Despite of considerable freezing artefacts, the presence or absence of specific reactivity was clearly recognizable in all tested individuals. Scale bars = 40 µm.

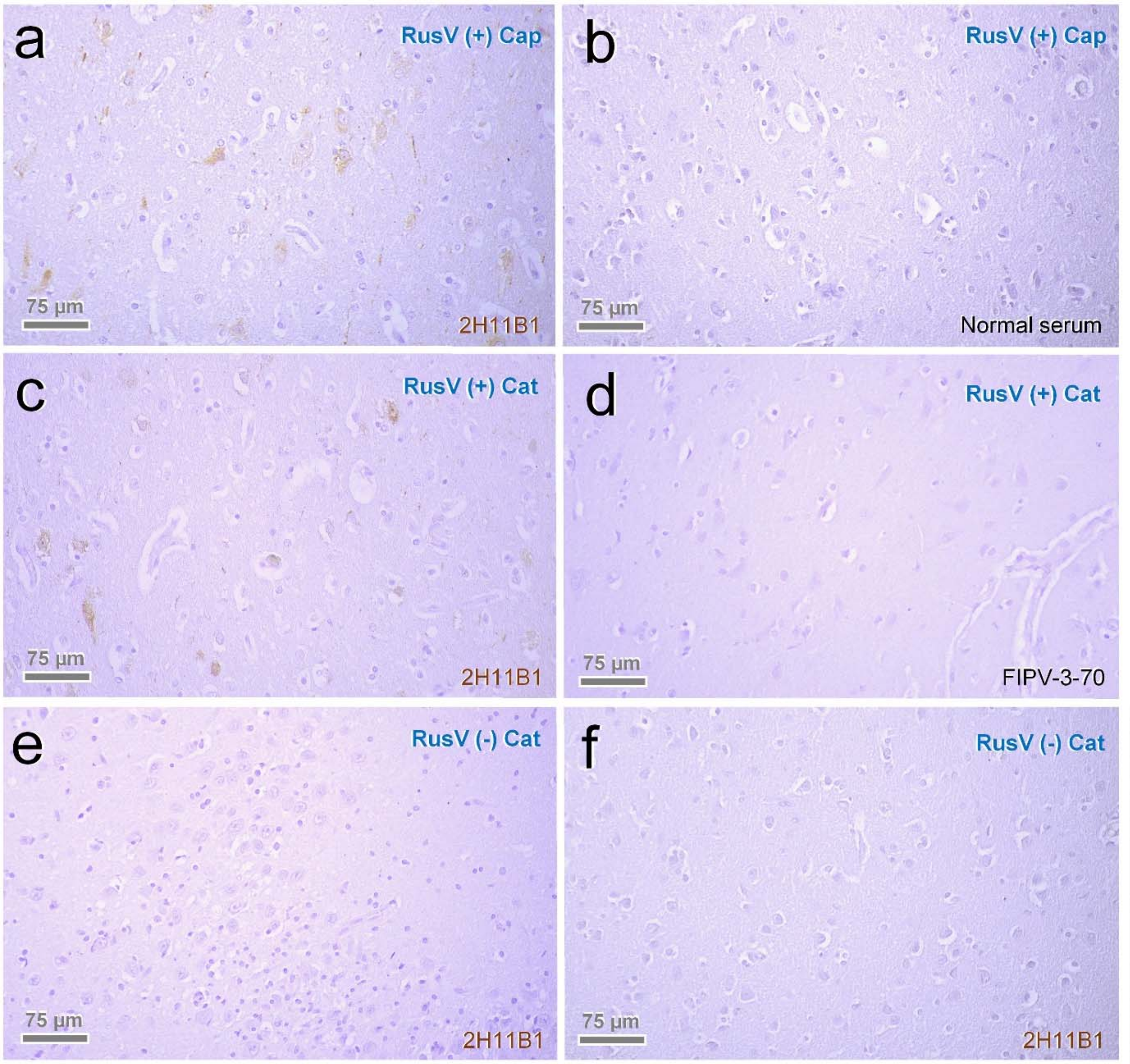


**Supplemental Figure S8: Control reactions of immunohistochemistry.** PCR-confirmed RusV-positive capybara (Cap) brain was used as positive control for RusV IHC using mouse monoclonal antibody 2H11B1 targeting the RusV capsid protein (**a**). Nonspecific staining effects were evaluated by replacement of 2H11B1 by normal serum (**b**) as well as by using another murine primary antibody in RusV-positive cats (**c, d**). Also brains of PCR-negative encephalitic (**e**) and non-encephalitic (**f**) controls underwent RusV IHC. (**a-f**) Counterstain: hematoxylin.

**Supplemental Table S1. Primers and probes used for bornavirus and rustrela virus (RusV) RNA detection, internal control RNA amplification, and Sanger sequencing.**

| Assay | Primer/Probe name | Sequence (5' to 3') | Reference |
|---|---|---|---|
| panBorna v7.2 | Borna-1319-F | CGCGACCMTCGAGYCTRGT | Schlottau et al.[5] |
| | Borna-1471.2-FAM | FAM-AAGAAYCCHTCCATGATCTCMGAYCMAGA-BHQ1 | Schlottau et al.[5] |
| | Borna-1529-R | GACARCTGYTCCCTTCCKGT | Schlottau et al.[5] |
| BoDV-1 Mix-1 | BoDV-1_1258+ | TAGTYAGGAGGCTCAATGGCA | Schlottau et al.[5] |
| | BoDV-1_1316_FAM | FAM-AAGAAGATCCCCAGACACTACGACG-BHQ1 | Schlottau et al.[5] |
| | BoDV-1_1419- | GTCCYTCAGGAGCTGGTC | Schlottau et al.[5] |
| RusV Mix-1 | RusV_1072+ | CGAGCGTGTCTACAAGTTCA | Bennett et al.[1] |
| | RusV_1161_P | FAM-CCGAGGAGGACGCCCTGTGC-BHQ1 | Bennett et al.[1] |
| | RusV_1237- | GACCATGATGTTGGCGAGG | Bennett et al.[1] |
| panRusV-2 | RusV_234+ | CCCCGTGTTCCTAGGCAC | this study |
| | RusV_256_P | TCGCCCCATTCWACCCAATT | this study |
| | RusV_323- | FAM-GTGAGCGACCACCCAGCACTCCA-BHQ1 | this study |
| eGFP mix 1 | EGFP-1-F | GACCACTACCAGCAGAACAC | Hoffmann et al.[6] |
| | EGFP-Probe1_HEX | HEX-AGCACCCAGTCCGCCCTGAGCA-BHQ1 | Hoffmann et al.[6] |
| | EGFP-2-R | GAACTCCAGCAGGACCATG | Hoffmann et al.[6] |
| conventional RusV RT-PCR & Sanger sequencing | RusV_80+ | GTCGAGGAGCAGATAAGCCC | this study |
| | RusV_528- | AGCGCCGGGTCYGTRACAAC | this study |

**Supplemental Table S2. Detection of rustrela virus (RusV) RNA and antigen in cats with 'staggering disease'.**

| Cat ID | CNS tissue used for RNA extraction | RusV RNA and antigen detection | | | | | GenBank accession |
|---|---|---|---|---|---|---|---|
| | | total [a] | HTS | RT-qPCR (Cq value) | ISH (score) [b] | IHC (score) [b] | |
| SWE_01 | FFPE | pos | *n.d.* | 35.1 | 1 | 2 | |
| SWE_02 | FFPE | pos | *n.d.* | 28.9 | 2 | 1 | ON641056 |
| SWE_03 | FFPE | pos | *n.d.* | 29.5 | 3 | 2 | ON641057 |
| SWE_04 | FFPE | pos | *n.d.* | 30.0 | 2 | 1 | ON641058 |
| SWE_05 | FFPE | pos | *n.d.* | 29.5 | 3 | 2 | ON641059 |
| SWE_06 | FFPE | pos | pos | 27.4 | 1 | 1 | ON641060 |
| SWE_07 | FFPE | pos | pos | 28.8 | 1 | 1 | ON641061 |
| SWE_08 | FFPE | pos | pos | 31.6 | 2 | 2 | ON641062 |
| SWE_09 | FFPE | pos | pos | 28.1 | 2 | 2 | ON641063 |
| SWE_10 | FFPE | pos | pos | 27.5 | 2 | 3 | ON641064 |
| SWE_11 | FFPE | pos | pos | 28.5 | 3 | 1 | ON641065 |
| SWE_12 | FFPE | pos | *n.d.* | 29.6 | 2 | 1 | ON641066 |
| SWE_13 | frozen | pos | pos | 23.7 | *n.d.* | 2 | ON641044 |
| SWE_14 | frozen | pos | pos | 20.4 | 3 | 2 | ON641045 |
| SWE_15 | frozen | pos | pos | 20.2 | 3 | 3 | ON641046 |
| AUT_01 | FFPE | pos | *n.d.* | 36.2 | 1 | 1 | |
| AUT_02 | frozen | pos | pos | 24.4 | 1 | 1 | ON641041 |
| AUT_03 | FFPE | pos | *n.d.* | neg | neg | 1 | |
| AUT_04 | FFPE | pos | neg | 34.0 | 2 | 3 | |
| AUT_05 | frozen | pos | *n.d.* | 30.7 | uncertain | neg | ON641050 |
| AUT_06 | frozen | pos | pos | 23.0 | 2 | 1 | ON641042 |
| AUT_07 | frozen | pos | *n.d.* | 28.5 | 1 | 1 | ON641051 |
| AUT_08 | frozen | pos | pos | 26.8 | 1 | 1 | ON641052 |
| AUT_09 | frozen | pos | *n.d.* | 29.4 | 2 | 3 | ON641053 |
| GER_01 | FFPE | pos | pos | 32.9 | uncertain | 1 | ON641055 |
| GER_02 | FFPE | pos | *n.d.* | neg | *n.d.* | 1 | |
| GER_03 | frozen | neg | *n.d.* | neg | *n.d.* | neg | |
| GER_04 | frozen | pos | pos | 28.9 | 1 | 1 | ON641043 |
| GER_05 | frozen | pos | *n.d.* | 31.6 | neg | 2 | ON641054 |
| Positive | | 28/29 | 14/15 | 26/29 | 22/26 | 27/29 | |

CNS: central nervous system; FFPE: formalin-fixed paraffin-embedded; HTS: high throughput sequencing followed by metagenomic analysis; Cq: cycle of quantification; ISH: *in-situ* hybridization using RNAscope; IHC: immunohistochemistry; pos: positive; neg: negative; n.d.: not determined

a Cats were considered RusV-positive if RusV RNA or antigen was detected by at least one of the applied methods.

b The scoring of IHC and RNAscope ISH signals is described in Supplemental Table S5.

**Supplemental Table S3. Demographic data on encephalitic cats included in this study.**

| Cat ID | Cat ID (submitter) | Month of death | Disease duration (weeks) | Location of origin | Age (years) | Sex [a] | Outdoor access |
|---|---|---|---|---|---|---|---|
| SWE_01 | O10/17 | 01-2017 | 1 | Skutskär | 6.8 | MN | yes |
| SWE_02 | O21/17 | 01-2017 | 3.7 | Rimbo | 1.5 | MN | yes |
| SWE_03 | O36/17 | 02-2017 | 2 | Bålsta | 7.7 | MN | yes |
| SWE_04 | O37/17 | 02-2017 | 1 | Uppsala | 6.1 | FN | yes |
| SWE_05 | O201/17 | 05-2017 | 1.9 | Krylbo | 3.1 | MN | yes |
| SWE_06 | O202/17 | 05-2017 | 0.7 | Gävle | 7.9 | MN | yes |
| SWE_07 | O313/18 | 10-2018 | 2 | Tärnsjö | 6.5 | MN | yes |
| SWE_08 | O13/19 | 02-2019 | 7 | Östervåla | 3.0 | FN | yes |
| SWE_09 | O96/19 | 03-2019 | 1.9 | Alunda | 1.9 | MN | yes |
| SWE_10 | O167/19 | 05-2019 | 1.1 | Uppsala | 12.3 | MN | yes |
| SWE_11 | PAT 6729/19 | 07-2019 | 6 | Björklinge | 7.5 | MN | yes |
| SWE_12 | PAT 6755/19 | 07-2019 | 8 | Uppsala | 3.5 | MN | yes |
| SWE_13 | 21_092-17 | 04-2021 | 58 | Stockholm | 7.0 | FN | yes |
| SWE_14 | O305/21 | 11-2021 | 1 | Vattholma | 3.3 | MN | yes |
| SWE_15 | O327/21 | 12-2021 | 0.6 | Uppsala | 2.8 | MN | yes |
| AUT_01 [c] | 1961/91 | 10-1991 | 1 | Ollersdorf | 2.0 | FN | yes |
| AUT_02 [c] | 706/92 | 04-1992 | N/A | N/A | adult | MN | N/A [b] |
| AUT_03 [c] | 1230/92 | 07-1992 | 4 | Vienna | 1.5 | MN | yes |
| AUT_04 [c] | 548/93 | 03-1993 | 8 | Gänserndorf | 2.0 | MN | yes |
| AUT_05 [c] | 1009/93 | 06-1993 | 3 | Glinzendorf | 2.0 | MN | yes |
| AUT_06 [c] | 1533/93 | 08-1993 | 1 | Prottes | 1.5 | MN | yes |
| AUT_07 [c] | 1807/93 | 10-1993 | 4 | Obersiebenbrunn | 2.0 | M | yes |
| AUT_08 [c] | 1812/93 | 10-1993 | 1 | Vienna | 1.5 | M | yes |
| AUT_09 [c] | 2281/93 | 12-1993 | N/A | Gänserndorf | 1.5 | M | yes |
| GER_01 [d] | S426/17 | 05-2017 | 2 | Hannover | 2.1 | MN | yes |
| GER_02 | NP 387/19 | 07-2019 | 16 | Leipzig | 5.0 | MN | yes |
| GER_03 | 21TRD0484 | 03-2021 | 0.1 | Stralsund | 13.0 | FN | N/A |
| GER_04 | 21TRD0953 | 07-2021 | 5 | Usedom | 3.0 | FN | yes |
| GER_05 | S34/22 | 01-2022 | 0.3 | Berlin | 6.0 | F | yes |

a F: female; M: male; N: neutered

b N/A: information not available

c Described in detail in previous publications[7, 8].

d Described in detail as case 1 in Nessler et al.[9].

**Supplemental Table S4. Summary of clinical signs, outcome, and histopathological lesions in rustrela virus (RusV)-infected cats**

| Cat ID | Clinical signs and outcome | Histological findings |
|---|---|---|
| SWE_01 | Ataxia, stiff gait, progressive inability to walk and stand, inability to retract claws; reduced/loss of menace response, palpebral, pupillary reflexes, postural (especially front limbs) reactions, withdrawal, panniculus, and perineal reflexes; depression, intermittent panting, obtundation; euthanasia after disease duration of one week | Mild multifocal lymphohistiocytic MEM [a] |
| SWE_02 | Ataxia, slow gait, falling over; reduced postural reactions (especially hind limbs); mild anisocoria, increased vocalization, contact seeking, hyperthermia, hyporexia, weight loss, pneumonia, obtundation; euthanasia after disease duration of 26 days | Moderate multifocal lymphohistiocytic MEM |
| SWE_03 | Ataxia, falling over; reduced menace response and postural reactions; euthanasia after approximately two weeks of disease duration | Moderate multifocal lymphohistiocytic MEM |
| SWE_04 | Ataxia, weakness of front limbs; reduced/loss of menace response, palpebral reflex, postural reactions, and withdrawal reflexes of front limbs; tremor, hyperesthesia at lumbar spine, mild anisocoria, obtundation, hyperthermia, hyporexia; euthanasia after disease duration of six days | Severe multifocal lymphohistiocytic MEM |
| SWE_05 | Ataxia, falling over, weakness, hypermetria, stiff front limbs on flexion, increased muscle tone of all limbs, tremor; reduced/loss of menace response, palpebral reflex, postural reaction, and withdrawal reflexes; increased vocalization, nervous, staring gaze; euthanasia after disease duration of 13 days | Mild to moderate multifocal lymphohistiocytic MEM |
| SWE_06 | Ataxia, falling over; reduced/loss of menace response and postural reactions (especially hind limbs); fecal incontinence, withdrawn behaviour, somnolence, obtundation; euthanasia after disease duration of five days | Mild to moderate multifocal lymphohistiocytic MEM |
| SWE_07 | Ataxia, wide-legged gait (especially hind limbs), difficulty in jumping, inability to retract claws; reduced panniculus and perianal reflexes; hyperesthesia at lumbar area and tail, kyphosis, hyperthermia, hyporexia, obtundation, cystitis; euthanasia after disease duration of approximately two weeks | Severe multifocal lymphohistiocytic MEM |
| SWE_08 | Ataxia/paresis, weakness of hind limbs, difficulty in jumping, inability to retract claws at hind limbs, hyperesthesia at lumbar back, pelvis, tail and caudal abdomen; increased vocalization, withdrawn behaviour, obtundation, initial hyperthermia, hyporexia, weight loss, pollakisuria, stranguria; euthanasia after disease duration of approximately seven weeks | Severe multifocal lymphohistiocytic MEM |
| SWE_09 | Ataxia, stiff gait, difficulty in jumping; loss of menace response; increased vocalization, contact seeking, hyperthermia, hyporexia, weight loss; euthanasia after disease duration of 13 days | Severe multifocal lymphohistiocytic MEM |
| SWE_10 | Ataxia, stiff gait, intermittent rigidity of right limbs; reduced/loss of menace response, palpebral reflex, and postural reflexes; disorientation, staring gaze, obtundation, hyporexia, weight loss; euthanasia after disease duration of eight days | Moderate multifocal lymphohistiocytic MEM |

(continue on next page)

(continued from previous page)

| | | |
|---|---|---|
| SWE_11 | Ataxia, increased muscle tone, left head tilt, progressive lateralized signs, tremor, generalized seizures; euthanasia after disease duration of approximately six weeks | Mild multifocal lymphohistiocytic MEM |
| SWE_12 | Ataxia, difficulties in jumping, falling over, paresis, weakness, somnolence, difficulties in drinking; euthanasia after disease duration of approximately eight weeks | Moderate multifocal lymphohistiocytic MEM |
| SWE_13 | Ataxia, increased muscle tone, tetraparesis; reduced/loss of menace response and postural reactions; depression; euthanasia after disease duration of 58 weeks | Moderate multifocal lymphohistiocytic MEM |
| SWE_14 | Ataxia, paresis, increased muscle tone, recumbency; reduced/loss of cranial nerve reflexes; increased salivation, obtundation; euthanasia after disease duration of seven days | Moderate multifocal lymphohistiocytic MEM |
| SWE_15 | Ataxia, wide-legged gait, fasciculations, behavioural changes, increased vocalization; euthanasia after disease duration of four days | Severe multifocal lymphohistiocytic meningoencephalitis [b] |
| AUT_01 [c] | Ataxia of one hind limb progressing to hemiparesis; euthanasia after disease duration of one week | Severe multifocal lymphohistiocytic MEM |
| AUT_02 [c] | Paralysis (no further records) | Moderate multifocal lymphohistiocytic MEM |
| AUT_03 [c] | Ataxia of hind limbs leading to flaccid paralysis; obtundation, hyporexia, hyperthermia; euthanasia after disease duration of four weeks | Severe multifocal lymphohistiocytic MEM |
| AUT_04 [c] | Ataxia of hind limbs, inability to jump and walk stairs, inability to retract claws; mydriasis, somnolence; euthanasia after disease duration of eight weeks | Severe multifocal lymphohistiocytic MEM |
| AUT_05 [c] | Ataxia and paralysis of hind limbs, circling; inability to retract claws, inappetence, somnolence, recumbency, hyperthermia; euthanasia after disease duration of three weeks | Moderate multifocal lymphohistiocytic MEM |
| AUT_06 [c] | Ataxia, falling over, recumbency with spastic tetraparesis, opisthotonus; euthanasia after disease duration of one week | Mild to moderate multifocal lymphohistiocytic MEM |
| AUT_07 [c] | Stiff gait, inability to jump, falling over, affectionate behaviour; euthanasia after disease duration of four weeks | Moderate multifocal lymphohistiocytic MEM |
| AUT_08 [c] | Ataxia, spastic paresis of hind limbs; euthanasia after disease duration of one week | Moderate multifocal lymphohistiocytic MEM |
| AUT_09 [c] | Stiff gait, inability to jump, falling over; euthanasia (disease duration not recorded) | Severe multifocal lymphohistiocytic MEM |

(continue on next page)

(continued from previous page)

| | | |
|---|---|---|
| GER_01 [d] | Compulsive pacing, increased muscle tone, whole body tremor, hyperthermia, hyporexia, disorientation, obtundation; euthanasia after disease duration of two weeks | Moderate multifocal lympho-histiocytic and plasmacytic MEM and vasculitis |
| GER_02 | Reduced postural reactions of all four limbs; absent menace response on both eyes and absent physiological nystagmus, obtundation, disorientation; euthanasia after disease duration of 16 weeks | Moderate multifocal lymphohistiocytic MEM |
| GER_03 [e] | Ataxia, salivation, inability to drink, aggression; euthanasia after disease duration of one day | Multifocal mild to moderate lymphohistiocytic encephalitis [b] |
| GER_04 | Ataxia of hind limbs, aggression, hyperthermia, apathy, hyporexia; euthanasia after disease duration of five weeks | Multifocal severe lymphohistiocytic MEM |
| GER_05 | Stupor, spasm, behavioural changes (aggression); euthanasia after disease duration of two days | Moderate multifocal lymphohistiocytic menigoencephalitis [b] |

a MEM: meningoencephalomyelitis

b Spinal cord not available for histologic evaluation

c Described in detail in previous publications[7, 8].

d Described in detail as case 1 in Nessler et al.[9].

e Cat GER_03 tested negative for RusV infection in all employed assays.

**Supplemental Table S5. Scoring of RNAscope ISH and IHC signal in brain slices**

| Score | Description |
|---|---|
| negative | no specific staining detectable |
| uncertain | very few signals, not clearly associated with cellular structures |
| 1 | signals in the cytoplasm or processes of singular neurons or glial cells |
| 2 | signals in an average of 5-10 cells per field (in 40-fold magnification) |
| 3 | signals in an average of more than 10 cells (up to ~100) per field (in 40-fold magnification) |
